# Cas3 Mediated Target DNA Recognition and Cleavage is Independent of the Composition and Architecture of Cascade Surveillance Complex

**DOI:** 10.1101/666776

**Authors:** Siddharth Nimkar, B. Anand

## Abstract

In type I CRISPR-Cas system, Cas3 –a nuclease cum helicase– in cooperation with Cascade surveillance complex cleaves the target DNA. Unlike the Cascade/I-E, which is composed of five subunits, the Cascade/I-C is made of only three subunits lacking the CRISPR RNA processing enzyme Cas6, whose role is assumed by Cas5. How these differences in the composition and organisation of Cascade subunits in type I-C influences the Cas3/I-C binding and its target cleavage mechanism is poorly understood. Here, we show that Cas3/I-C is intrinsically a single-strand specific promiscuous nuclease. Apart from the helicase domain, a constellation of highly conserved residues –that are unique to type I-C– located in the uncharacterised C-terminal domain appears to influence the nuclease activity. Recruited by Cascade/I-C, the HD nuclease of Cas3/I-C nicks the single-stranded region of the nontarget strand and positions the helicase motor. Powered by ATP, the helicase motor reels in the target DNA, until it encounters the roadblock en route, which stimulates the HD nuclease. Remarkably, we show that Cas3/I-C supplants Cas3/I-E for CRISPR interference in type I-E *in vivo*, suggesting that the target cleavage mechanism is evolutionarily conserved between type I-C and type I-E despite the architectural difference exhibited by Cascade/I-C and Cascade/I-E.

## INTRODUCTION

Archaea and Bacteria defend themselves from the assault of phages and plasmids by employing a genetically encoded RNA mediated adaptive immunity referred to as CRISPR-Cas system. CRISPR constitutes an array of direct repetitive sequences (each of about 30 - 40 bp) that are intervened by similarly sized variable spacer sequences. These spacer sequences are preferentially captured from the invading foreign genetic element and get integrated into the CRISPR array. They serve as immunological memory - akin to antibodies in higher organisms - to specifically recognise and target the invader during recurrent infection (1–5). CRISPR based targeting can be operationally distinguished into three distinct stages: (i) Adaptation, (ii) Maturation and (iii) Interference. Immunological memory is acquired during adaptation wherein short fragment of DNA from the foreign genetic element (protospacer) is acquired and incorporated into the CRISPR locus in a polarised fashion (6–10). This is followed by the transcription and processing of the pre-CRISPR transcript that gives rise to the mature CRISPR RNA (crRNA) onto which either a single (Class II) or several Cas protein(s) (Class I) assemble(s) to form a ribonucleoprotein (RNP) surveillance complex. The crRNA within the RNP guides the target recognition by base complementarity whereas the protein component(s) facilitate(s) the cleavage of the target nucleic acid (1,3,4,11–13).

CRISPR-Cas system is highly diverse and is broadly divided into two classes (I & II), which are further divided into six types (I-VI) and several subtypes (14–17). In Class II system, the RNP surveillance complex is comprised of a single Cas protein *viz.*, Cas9/II-A (18,19) or Cpf1/V-A (20) and this effector complex participate in both target recognition and cleavage. On the contrary, the surveillance complex in Class I system is highly variable and comprises of several Cas proteins (3,21–23), which are involved in target recognition alone.

In Class I/type I system, the effector complex for CRISPR interference is referred to as Cascade – which comprises of crRNA, a large and a small Cas subunits, variable number of Cas5, Cas6, and Cas7 – that recognize the target and recruits Cas3 – a trans-acting motor protein – that cleaves the target DNA (21,24–26). Target identification by Cascade complex requires that the upstream of target sequence contains a short nucleotide sequence (2-4 nt) called proto-spacer adjacent motif (PAM), which is present only in the invading phages/plasmids (27). The primary function of the PAM sequence is to discriminate self versus non-self and thereby pre-empt targeting of the host genome (12,28). In line with this, mutations in the PAM sequence leads to target evasion (24,29,30). Target identification by Cascade leads to R-loop formation, wherein the target strand base pairs with the crRNA and the non-target strand becomes single-stranded (31–36). It appears that Cas3 recruitment is facilitated by a conformational change in the large subunit of Cascade complex – say Cas8E in type I-E – and this leads to Cas3 mediated preferential cleavage of the non-target strand (30,37–40). Cas3 acts on the invasive DNA by cooperative interaction between its “HD” nuclease and Superfamily 2 (SF2) “DEXD/H” helicase domains (41,42). The HD domain shows divalent metal ion-dependent nuclease activity with a preference for single-stranded DNA (43–45). The DEXD/H box helicase domain comprises of two juxtaposed RecA domains, and these are followed by a long linker and a C-terminal domain (CTD), whose function is poorly defined. After recruitment by Cascade, nuclease domain of Cas3 binds to non-target strand after making a single-stranded nick, followed by processive degradation of target DNA (35,46).

Befitting the enormous diversity exhibited by type I system, the composition and architecture of Cascade effector complex have undergone commensurate variation (14,15). In tune with this, the domain architecture of Cas3 in some sub-types has evolved in a very distinctive way such that HD and Helicase domains, which are fused in some sub-types, have undergone domain fission in others (44,47,48). The current mechanistic understanding on CRISPR interference in type I stems majorly from the prototypical type I-E system (24,30,33,36,39,41,42,46,49,50) with sporadic investigations from other type I systems such as I-A, I-F and I-G (47,48,51–58). In type I-C, the Cascade complex is composed of only three subunits, *viz*. Cas5, Cas7 and Cas8, as against five (CasA-E) in case of type I-E (23,25). Cas6 – which is essential for crRNA maturation – is absent in type I-C and Cas5, which is inert in type I-E, has supplanted the role of Cas6 (23,59,60). Given the architectural differences among Cascade complex of type I-C and type I-E, it is reasonable to assume that this will be translated into functional differences. However, apart from type I-E, studies that systematically investigated the function of Cas3, as well as the functional interaction between Cascade and Cas3 in type I-C, are virtually non-existent hitherto. Arguably, this is partly due to the challenges associated with the reconstitution of Cascade complex and Cas3 in type I-C *in vitro*. Here, we have attempted to elucidate the functional mechanism of CRISPR interference in type I-C both by reconstituting the Cascade/I-C and Cas3/I-C *in vitro* and *in vivo*. This revealed the unique features of type I-C interference machinery as well as the conserved functional mechanism with type I-E system.

## MATERIAL AND METHODS

### Strain construction

For studies related to the integration of genes encoding Cascade/I-C (Cas5, Cas8 and Cas7) from *B. halodurans, E. coli* IYB5101 (vide Supplementary Table S2) was used as a recipient strain. This strain harbours T7 polymerase, which is under the control of *ara*C and hence it is inducible with arabinose. Genes encoding Cascade/I-C (Cas5, Cas8 and Cas7) were amplified as a polycistronic construct using *B. halodurans* C-125 genomic DNA as a template. This construct has T7 promoter sequence upstream of the coding region. Subsequently, this construct was integrated into *E. coli* IYB5101 using pOSIP-CT (addgene #45981) to create the strain IC-I through one-step cloning and chromosomal integration of DNA (clonetegration), which uses phage integrases (61). Double knock-out of *cas3* (Δcas3) and *hns* (Δhns) was created in *E. coli* K-12 BW25113 (vide Supplementary Table S2) using λ Red recombineering (62) to create strain IC-2. The list of strains employed for this study is presented in Supplementary Table S2.

### Molecular Cloning

*Bacillus halodurans* C-125 strain was procured from The Microbial Type Culture Collection and Gene Bank (MTCC), India. Gene encoding Cas3 (BH0336) and all its mutants were cloned in 1R (addgene #29664) at SspI restriction site by employing Gibson Assembly. Point mutations were created in Cas3/I-C using PCR based mutagenesis. The 1R vector encodes Strep II tagged protein. Genes encoding Cascade/I-C (Cas5, Cas8 and Cas7) were amplified as a polycistronic construct and inserted into the 1R expression vector to produce pCascade/I-C. CRISPR array containing seven copies of identical repeat-spacer units that were preceded and succeeded by T7 promoter and terminator sequences, respectively, was commercially synthesized (GenScript) and inserted into 13S-R (addgene # 48328) to produce pCRISPR/I-C. 13S-R and 1R are compatible vectors and can be used for purification of Cascade/I-C complex. Target DNA sequence was inserted into the pUC19 vector using KpnI and HindIII restriction sites. All cloned constructs were verified by Sanger sequencing. The information on the oligonucleotides used in this study can be found in Supplementary Table S1.

### Protein purification

*E. coli* BL21 (DE3) harbouring pWT-1 (Cas3/I-C cloned in 1R) was grown in LB broth supplemented with 50 μg/ml kanamycin at 37 °C till the OD_600_ reached 0.6. At this point, Cas3/I-C expression was induced with 0.2 mM IPTG and cells were allowed to grow for 12 hrs at 18 °C. Cells were harvested and re-suspended in binding buffer (20 mM Tris-Cl (pH 8.0), 300 mM NaCl, 10 % glycerol, 1 mM phenylmethylsulfonyl fluoride (PMSF) and 6 mM β-mercaptoethanol). Cells were lysed using cell disruptor (20 kpsi), and cell debris was removed by centrifugation at 4 °C and 16,000 x g for 30 mins. After cell lysis, DNase I (50 μg/ml) was added to reduce the viscosity due to the presence of intact genomic DNA. The clear supernatant was loaded onto pre-equilibrated 5ml StrepTrap HP (GE Healthcare) column. After loading, the column was washed with 10 CV of binding buffer to remove unbound protein and Cas3/I-C was eluted in binding buffer containing 2.5 mM desthiobiotin. Cas3/I-C was further loaded onto HiTrap Heparin HP column (GE Healthcare) to remove impurities. Subsequently, it was eluted with a linear gradient of 0.15 to 2 M NaCl in binding buffer. Later, the protein was further purified through HiLoad 16/600 Superdex 200 prep grade column (GE Healthcare) and eluted in buffer containing 20 mM Tris-Cl (pH 7.5), 150 mM NaCl, 10 % glycerol, and 6 mM β-mercaptoethanol. After concentration, samples were flash frozen in liquid nitrogen and stored at −80 °C until use. Protein samples were run on SDS-PAGE to check the purity and concentration was measured by absorbance at 280 nm. Cas3/I-C mutants cloned in 1R (pD48A, pQ253A, pD395A, pK742A, pK743A, pQ745A, pQ746A and pY747A) were purified using similar protocol mentioned above.

*E. coli* BL21 (DE3) harbouring pCascade/I-C and pCRISPR/I-C (vide Supplementary Table S3) was grown in LB broth supplemented with 50 μg/ml kanamycin and 100 μg/ml spectinomycin at 37 °C till the OD_600_ was equal to 0.6. Cascade/I-C expression was induced with 0.2 mM IPTG and cells were allowed to grow overnight at 25 °C. Cells were harvested and re-suspended in binding buffer (20 mM Tris-Cl (pH 7.5), 150 mM NaCl, 10 % glycerol, 1 mM PMSF, and 6 mM β-mercaptoethanol). Cells were lysed using cell disruptor (20 kpsi) and cell debris was removed by centrifugation at 4 °C and 16,000 x g for 30 mins. After cell lysis, DNase I (50 μg/ml) was added to reduce the viscosity. The clarified supernatant was passed through pre-equilibrated 5ml StrepTrap HP (GE Healthcare) column. After washing with 10 CV of binding buffer, Cascade/I-C was eluted in binding buffer containing 2.5 mM desthiobiotin. Samples were pooled up and passed through HiTrap Heparin HP column (GE Healthcare) and eluted with a linear gradient of 0.15 to 2 M NaCl in the binding buffer to remove additional impurities. Later, proteins were further purified through HiLoad 16/600 Superdex 200 prep grade column (GE Healthcare) and eluted in buffer containing 20 mM Tris-Cl (pH 7.5), 150 mM NaCl, 10 % glycerol, and 6 mM β-mercaptoethanol. Concentrated samples were flash frozen in liquid nitrogen and stored at −80 °C.

### ATP Hydrolysis Assay

ATP hydrolysis reactions were conducted at 37 °C in the reaction buffer containing 20 mM Tris-Cl (pH 8.0), 50 mM NaCl, 10 mM Mg^2+^, 1 mM ATP, 6 mM β-Mercaptoethanol, and 1 μM Cas3. Hydrolysis reactions were performed in the presence of 10 ng of ssDNA, dsDNA and 16S rRNA. Malachite Green assay was used to determine the liberated inorganic phosphate. Reactions were initiated by addition of ATP and stopped by adding malachite green reaction mixture (ammonium molybdate: malachite green in ratio 1:4). To this mixture, 34 % sodium citrate was added to make a final concentration of 3.7 %. The volume was made up to 1 ml using water and absorbance was measured at 630 nm. A standard curve between absorbance and inorganic phosphate concentration was established by using varying concentration of KH_2_PO_4_. All the reactions were performed in triplicates.

### Electrophoretic mobility shift assay

Single-stranded DNA constructs (Supplementary Table S1) with or without 6-FAM labels were purchased from Integrated DNA Technologies, Inc (IDT) and gel purified to remove truncated DNA fragments. A 100 bp target DNA construct was generated by annealing the oligonucleotides having 3’ complementarity and subjecting them to extension in a PCR using Pfu DNA polymerase. 50 ng of target DNA was incubated with 500 nM of Cascade/I-C complex in the buffer containing 20 mM Tris-Cl (pH 8.0), 150 mM NaCl and 1 mM DTT at 37 °C for 30-60 mins to allow R-loop formation. Post incubation, one set of the sample was treated with 1 mg/ml proteinase K to test the release of target DNA bound to Cascade/I-C. Samples were directly loaded onto 20 % (w/v) native polyacrylamide gel and electrophoresed in 1X TBE at 4°C. While 6-FAM labelled constructs were directly visualized under UV in gel documentation system (Bio-Rad), unlabelled constructs were visualized after staining with ethidium bromide (EtBr).

### Assay for Nuclease Activity

In order to assess the nuclease activity of Cas3/I-C in the presence of Cascade/I-C, target DNA and Cascade/I-C were incubated with a varying concentration of Cas3/I-C in the buffer containing 20 mM Tris-Cl (pH 8.0), 60 mM NaCl, 1 mM dithiothreitol (DTT), 10 mM MgCl_2_ and 1 mM ATP. After the incubation, samples containing unlabelled DNA were directly loaded onto 20 % (w/v) native polyacrylamide gel and electrophoresed in 1X TBE at 4 °C. DNA bands were visualized in the gel documentation system (Bio-Rad) after staining with EtBr. 6-FAM labelled DNA samples were analysed on 20% (w/v) denaturing PAGE containing 6 M urea and directly visualised without any post-staining.

All other nuclease assays in the absence of Cascade/I-C were performed in cleavage buffer containing 20 mM Tris-Cl (pH 8.0), 60 mM NaCl, 1 mM dithiothreitol (DTT) and 10 mM MgCl_2_, unless specified otherwise. To check metal ion dependency, 2 mM of various metal salts and 500 nM Cas3/I-C were used. DNA substrates were used at 100 ng, whereas RNA substrates were used at 20 ng. The reaction was carried out at 37 °C for 60 mins. Cleavage products were visualised by either 0.8 % agarose gel electrophoresis or 10-15% (w/v) PAGE.

### Nuclease activity on bubble DNA construct

Unlabelled single-stranded DNAs (100 nt) having complementary terminal ends were purchased from IDT and PAGE purified to remove truncated DNA fragments (Dloop-AAG-FP and Dloop-TTC-RP; refer Supplementary Table S1). These oligonucleotides (1 μM) were annealed and incubated with increasing concentration of Cas3/I-C (0.2 to 1 μM) in the reaction mixture containing 20 mM Tris-Cl (pH 8.0), 150 mM NaCl, 1 mM DTT, 10 mM MgCl_2_ and 1 mM ATP at 37 °C for 120 mins. Samples were directly loaded onto 20 % (w/v) native polyacrylamide gel and electrophoresed in 1X TBE at 4 °C. Gels were post-stained with ethidium bromide (EtBr) and DNA bands were visualized in the gel documentation system.

### Nuclease activity on biotin labelled DNA construct

Biotin labelled DNA construct with 5’ 6-FAM label was designed and purchased from Bioserve Biotechnologies (India) Pvt. Ltd. and PAGE-purified to remove free 6-FAM and truncated oligonucleotides (ssDNA-biotin; refer Supplementary Table S1). To form double-stranded target DNA, both the strands were annealed by heating at 95 °C followed by gradual cooling. dsDNA or ssDNA was pre-incubated with streptavidin to form a roadblock in the molar ratio 2:3 (DNA: Streptavidin). Wherever mentioned, 500 nM of Cascade/I-C was added to form r-loop. Subsequently, Cas3/I-C was added in the reaction mixture containing 20 mM Tris-Cl (pH 8.0), 150 mM NaCl, 1 mM DTT, 10 mM MgCl_2_ and 1 mM ATP/ADP/AMP-PNP. Samples were analysed on 20 % (w/v) denaturing PAGE containing 8 M urea and directly visualised in the gel documentation system.

### Assay for Helicase activity

Partial DNA duplexes were used to test the helicase activity of Cas3/I-C and its variants. 6-FAM labelled oligonucleotides were obtained from IDT. To generate the construct with 3’ overhang, 3’ 6-FAM labelled short oligonucleotide (36 nt) and unlabelled long oligonucleotide (70 nt) were annealed (refer Supplementary Table S1). Similarly, in order to generate the construct with 5’ overhang, 5’ 6-FAM labelled short oligonucleotide (34 nt) and unlabelled long oligonucleotide (70 nt) were annealed (refer Supplementary Table S1). In addition, an excess of an unlabelled short oligonucleotide having an identical sequence as that of 6-FAM labelled was used as trap DNA, in order to avoid the re-annealing of the separated strand. Helicase activity was assessed by mixing Cas3/I-C (1 μM) and 6-FAM labelled DNA substrates (1 μM) in the reaction buffer containing 20 mM Tris-Cl, 60 mM NaCl, 6 mM β-mercaptoethanol, 10 mM MgCl_2_, 50 μM Trap-DNA and 1 mM ATP (or other triphosphate variants). The reaction was carried out at 37 °C for 2 hours. DNA bands were visualized on 12 % (w/v) native PAGE.

### Assay for CRISPR Interference

*E. coli* IC-1 harbouring pWT-1 and pCRISPR/I-C (vide Supplementary Table S3) was grown in LB broth supplemented with 25 μg/ml kanamycin, 50 μg/ml spectinomycin, 25 μg/ml chloramphenicol, 0.2 % L-arabinose and 0.1 mM IPTG at 37 °C until OD_600_ reached 0.3. Cells were harvested at 4 °C by centrifuging at 2700 x g. Harvested cells were made chemically competent using 0.1 M calcium chloride and transformed with pT1/pNT1 (see Supplementary Table S3). Cells were grown overnight at 37 °C on LB agar supplemented with 25 μg/ml kanamycin, 50 μg/ml spectinomycin, 25 μg/ml chloramphenicol, 50 μg/ml ampicillin, 0.2 % L-arabinose and 0.1 mM IPTG. The number of colonies obtained were counted and transformation efficiency was calculated using the equation mentioned below.

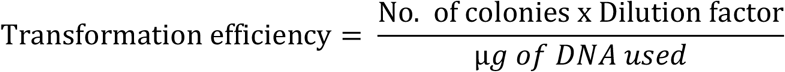

Transformation efficiency for Cas3 variants (pHD, pHL, pCas3ΔCTD, pK742A, pK743A, pQ745A, pQ746A and pY747A; vide Supplementary Table S3) was calculated using similar protocol.

*E. coli* IC-2 harbouring pWT-2 was cultured in LB broth supplemented with 50 μg/ml ampicillin and 20 μM IPTG at 37 °C. Cells were harvested when OD_600_ reached 0.3. Subsequently, harvested cells were made chemically competent as mentioned above. These cells were transformed with pRSF-11/pST-KT (see Supplementary Table S3). Cells were grown overnight in LB agar with 25 μg/ml kanamycin and 50 μg/ml ampicillin. Similarly, transformation efficiencies for other variants of Cas3 (pD48A, p253A, pD395A, pHD + pHL, pHD + pHLΔCTD) were determined.

### Fluorescence quenching assay

DNA strand with 5’ biotin and 6-FAM at 5^th^ nucleotide (NTS) and Iowa Black FQ (IB) at 29^th^ nucleotide was ordered from IDT. All the DNA constructs were PAGE purified to remove truncated fragments and annealed by heating at 95 °C followed by gradual cooling. 100 nM of annealed dsDNA was pre-incubated with 200 nM streptavidin and 500 nM Cascade/I-C. Cas3 at increasing concentration (0-5 μM) was added in the reaction mixture containing 20 mM Tris-Cl (pH 8.0), 150 mM NaCl, 1 mM DTT, 10 mM MgCl_2_ and 1 mM ATP/ADP/AMP-PNP and fluorescence intensity was measured immediately in FluoroMax^®^-4 (Horiba Scientific). For time-dependent measurement, 500 nM of Cas3/I-C was added in the reaction mentioned above and fluorescence intensity was measured for 120 min.

### Anisotropy measurements

Biotin labelled ssDNA construct with 5’ 6-FAM as mentioned previously (see Supplementary Table S1) was pre-incubated with streptavidin in the molar ratio 2:3 (DNA: Streptavidin). Cas3 was added in the reaction mixture containing 20 mM Tris-Cl (pH 8.0), 150 mM NaCl, 1 mM DTT, 10 mM MgCl_2_ and 1 mM ATP/ADP/AMP-PNP and anisotropy values were measured using FluoroMax^®^-4 (Horiba Scientific). To assess Cas3/I-C binding, anisotropy readings were taken immediately after adding Cas3 (0-10 μM). For time-dependent measurement of anisotropy, 500 nM of Cas3 was added and readings were noted for 90 min.

### Homology Modelling

A model of Cas3/I-C was prepared using I-TASSER server (63). Cas3/I-C (BH0336) sequence was uploaded on I-TASSER server and Cas3/I-C was modelled using Cas3/I-E from *Thermobifida fusca* [PDB ID: 4QQX and 6C66] as a template. Cas3/I-C from *B. halodurans* and Cas3/I-E from *T. fusca* share 38% sequence identity. The model with the highest C-score was chosen for further analysis.

## RESULTS

### CRISPR interference requires coordination between nuclease and helicase domains of Cas3/I-C

To test the substrate preference for HD nuclease, different nucleic acid substrates were utilised in the presence of ATP and Mg^2+^ ions (Figure 1 & Supplementary Figure S1A-F). This showed that Cas3/I-C cleaves both double-stranded (pQE2) and single-stranded (M13mp18) DNA proficiently (Figure 1B). Further, the nuclease activity does not differentiate between methylated (pQE2) and unmethylated (PCR amplicon) forms as substrates (Figure 1C). Prompted by the lack of apparent specificity for the target, we questioned whether Cas3/I-C shows any preference for the length of the target. We utilised DNA substrates of various lengths – 70 bp, 400 bp and 2.5 kb. We found that while Cas3/I-C is active against 2.5 kb substrate, it is barely active against 400 bp and 70 bp substrates, respectively (Figure 1D). In line with this, though it acts on long RNA, no nuclease activity was discernable with small RNA fragment (150 nt) (Figure 1E & Supplementary Figure S1). These suggest that despite the apparent preference for longer DNA, Cas3/I-C exists as a promiscuous nuclease with no apparent intrinsic specificity for the target.

**Figure 1.**
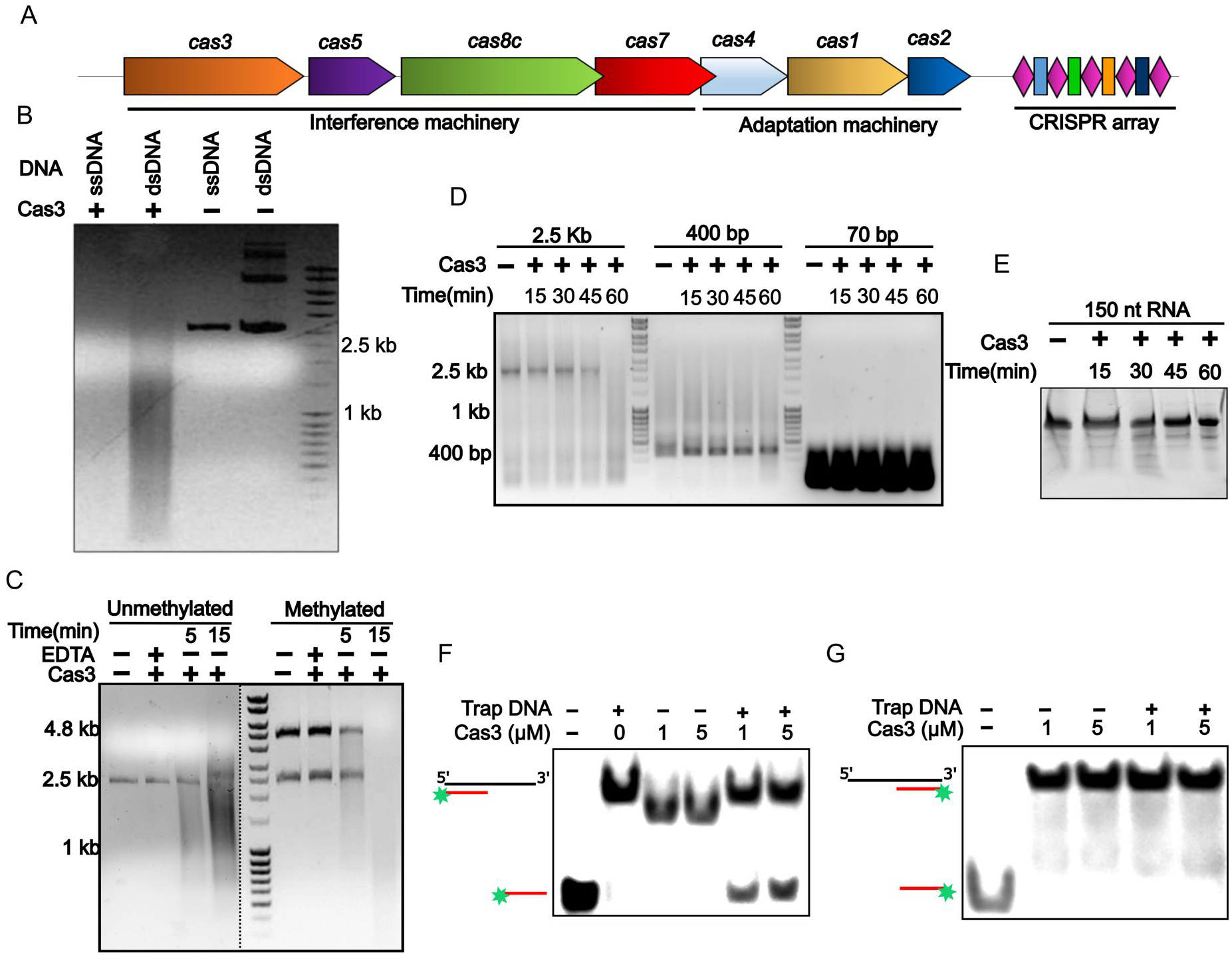
Cas3/I-C cleaves the target DNA indiscriminately. (A) A schematic overview of type I-C CRISPR-Cas locus from *B. halodurans* C-125. Genes are drawn to represent locus approximately to the scale. Effector nuclease (*cas3*), cascade (*cas5, cas8, and cas7*) and adaptation (*cas4, cas1, and cas2*) genes are shown. Repeats are shown as diamonds in magenta and spacers as rectangles in different colours. (B) ssDNA and dsDNA represent single-stranded and double-stranded DNA substrates, respectively. Cas3/I-C cleaves both single (M13mp18) and double (pQE2) stranded DNA in the presence of divalent metal ion (Mg^2+^) and ATP. (C) Methylated and unmethylated DNA fragments are of the same size (2.5 kb). Un-methylated DNA was generated using *Pfu* polymerase which generates blunt-end products. The methylated DNA fragment of 2.5 kb was released from plasmid using restriction enzymes NdeI and KpnI. An additional band of 4.8 kb corresponds to the linearised vector pQE2. DNase activity is not sensitive to methylation. The results were visualised using 0.8% agarose gel electrophoresis. Dotted line indicates discontinuity in the gel for clarity. (D-E) DNase/RNase activity depends on the size of the substrate. Cas3/I-C preferred large DNA substrates (2.5 kb), whereas smaller DNA (400 bp and 70 bp) and RNA (150nt) substrates were not completely cleaved even after 60 mins of incubation. DNase and RNase activities were checked on 0.8 % agarose and 12 % denaturing PAGE, respectively. (F-G) DNA unwinding was tested using fluorescently labelled partial DNA duplex. Displacement of 36 nt was observed when a substrate containing 3’ overhang was used. 15% native PAGE was used to analyse the results.

Having characterised the HD nuclease activity, we set out to assess the DEXD/H domain-mediated helicase activity using a non-specific DNA substrate. We utilised DNA containing 3’ and 5’ overhang substrates to assess the directionality of helicase activity (Figure 1F-G). We observed that Cas3/I-C unwinds only that DNA, which possesses a 3’ overhang, suggesting that it translocates in 3’-5’ direction in the presence of ATP and Mg^2^+ (Figure 1F-G & Supplementary Figure S2A-B). These suggest that Cas3/I-C is a generic 3’-5’ helicase with no apparent sequence specificity.

It is not clear whether the nuclease and helicase domains of Cas3/I-C cooperate during interference *in vivo*. To address this, we utilised *E. coli* IYB5101 as a surrogate host (IC-1, Supplementary table S2) for porting the type I-C system (CRISPR/I-C and Cascade/I-C) from *B. halodurans* (Figure 2A). We introduced the protospacer sequence abutting the targeting PAM sequence (TTC) into pUC19 vector, which acted as target plasmid (T1). On the other hand, an empty pUC19 vector lacking protospacer was used as the non-target plasmid (NT1). In the absence of Cas3/I-C, the transformation efficiency of both T1 and NT1 was comparable (Figure 2B). When Cas3/I-C was introduced via a plasmid-borne construct, the transformation efficiency of NT1 was retained; however, that of T1 was drastically reduced (Figure 2B). This suggests that the type I-C system is able to discriminate between the target and the non-target substrates *in vivo*. In addition, we observed that CRISPR interference is not active in those strains if Cas3/I-C lacks C-terminal (CTD) domain or possesses a mutation in HD or DEXD/H domain that inhibits nuclease and helicase activity, respectively (Figure 2B). However, co-expression of stand-alone HD and DExD/H domains resulted in functional assembly of interference complex as evidenced by reduction in transformation efficiency (Figure 2B). This suggests that the interplay between these domains is crucial for CRISPR interference in type I-C system.

**Figure 2.**
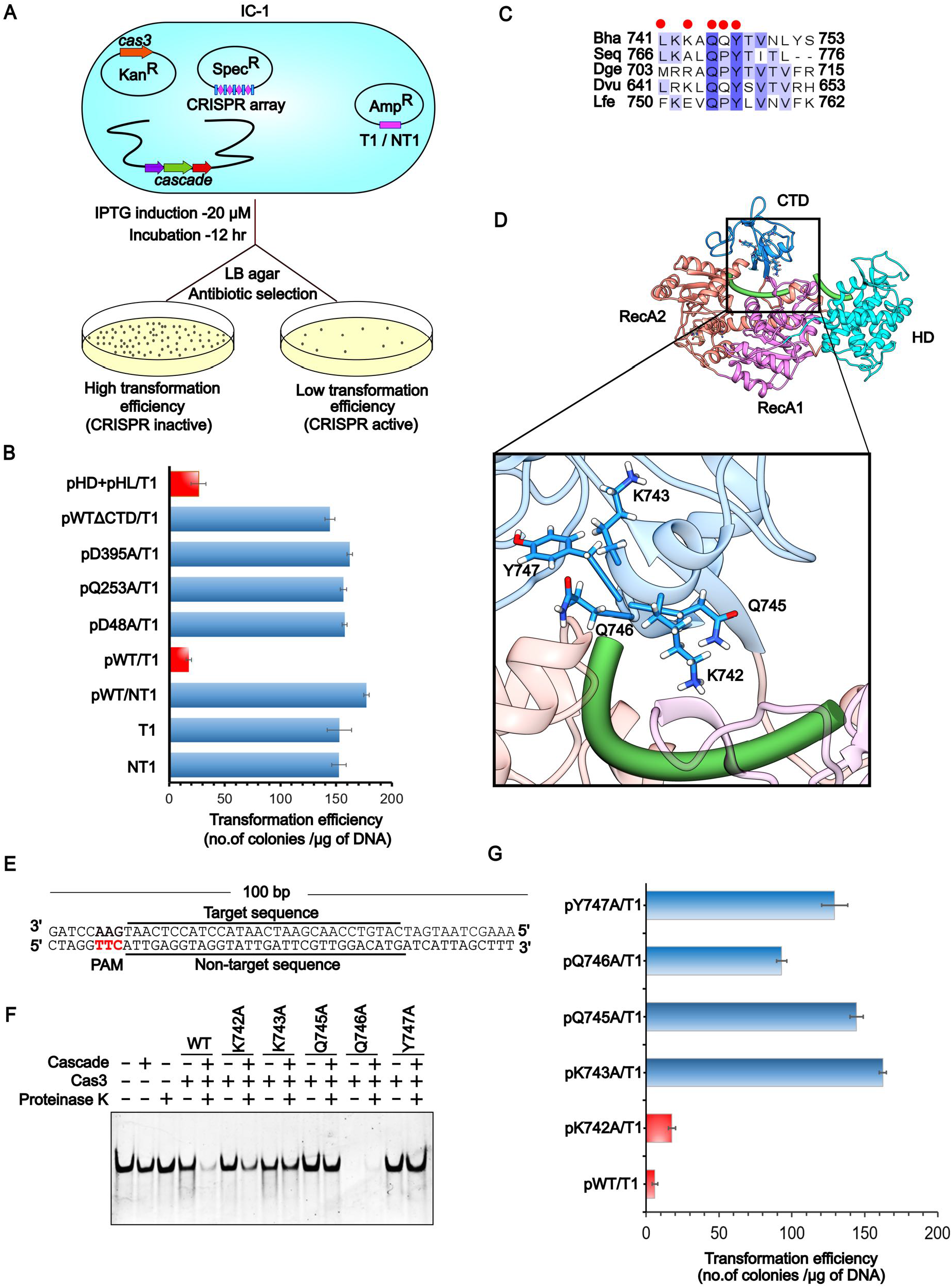
Inter-domain interaction in Cas3/I-C influences CRISPR interference. (A) An *E. coli* IYB5101 strain harbouring genes encoding Cascade/I-C (IC-1) was used as a surrogate for *in vivo* experiments. Cas3/I-C and crRNA were expressed through IPTG inducible vectors having kanamycin and spectinomycin antibiotic resistance markers, respectively. The target sequence was inserted into pUC19 vector (T1), whereas an empty pUC19 vector was used as non-target (NT1). Low transformation efficiency indicates functional CRISPR interference. (B) Transformation efficiencies of the strain IC-1 against the target (T1) and the non-target (NT1) for wild type and variants of Cas3/I-C are shown. In the presence of Cas3/I-C wild type (pWT) and coexpressed construct of HD and DExD/H domain (pHD+pHL) there was a significant reduction in transformation efficiency (indicated by a red bar). Interference was rendered ineffective when the HD domain mutant (pD48A), Helicase domain mutant (pQ253A and pD395A), and CTD deletion mutant (pWTΔCTD) were utilized. Error bar represents standard deviation measured from three independent trials. The boundary of the domain variants are as follows: HD domain (pHD) 1-248; DExD/H domain (pHL) 249-800; WT without CTD (pWTΔCTD) 1-709. (C) Highly conserved residues in the CTD domain from type I-C system are shown. Red dots indicate those residues from *B. halodurans* that are chosen for alanine-scanning mutagenesis. (D) Homology model of Cas3/I-C from *B. halodurans* based on Cas3/I-E structure [PDB ID: 4QQX] shows the organisation of nuclease (HD in cyan), DEXD/H helicase (RecA1 in pink and RecA2 in orange) and CTD (in blue) domains. DNA is shown as a rod in green. Amino acids (K742, K743, Q745, Q746, and Y747) from CTD domain that are possibly interacting with target DNA are shown. (E) A 100 bp target DNA harbouring functional PAM sequence TTC is depicted. Target sequence represents the strand, which is complementary to crRNA. (F) CTD point mutants were tested for their nuclease activity. Cascade/I-C was incubated with target DNA in the reaction mixture containing Mg^2+^ and ATP to form R-loop. Wild type or CTD mutant was added to the reaction and the cleavage was monitored. Proteinase K was added to release residual Cascade bound DNA fraction as observed using 20 % native PAGE. (G) CTD mutants were tested for their role in CRISPR interference using *E. coli* IC-1 strain as the surrogate host. Wild type and CTD mutants were expressed through an IPTG inducible 1R vector (referred to as pWT, pK742A, and so on). Transformation efficiency was measured against pUC19 harbouring the target sequence with functional PAM TTC (referred to as pT1). Error bar represents standard deviation measured from three independent trials.

### A constellation of highly conserved residues in the C-terminal domain of Cas3/I-C regulates target cleavage

Apart from the two major functional domains – nuclease and helicase – Cas3/I-C has a CTD, whose function is poorly defined (Figure 2D). Intrigued by the fact that CTD is crucial for CRISPR interference (Figure 2B) we set out to probe its role in more detail. Previously determined crystal structure of Cas3/I-E with target DNA [PDB ID: 4QQX] shows that CTD caps the two juxtaposed RecA domains (that are part of the DEXD/H domain) such that the target DNA is sandwiched between CTD and RecA domains (Figure 2D). Sequence alignment of Cas3/I-C orthologs showed high conservation of K742, Q745, and Y747 and these conserved residues were absent in Cas3/I-E, suggesting a functional significance for these residues in type I-C (Figure 2C & Supplementary Figure S3A). Since the structure of Cas3/I-C is not determined, we employed the Cas3/I-E structure [PDB ID: 4QQX] as template and made a homology model for Cas3/I-C, which suggested that K742, K743, Q745, Q746, and Y747 that are part of CTD could stabilize the interaction of the target DNA with the helicase core (Figure 2D). Based on this, we hypothesised that CTD may be involved in stabilising the interaction with target DNA so that Cas3/I-C does not dissociate from DNA during its translocation (Supplementary Figure S4). To test this hypothesis, we performed alanine-scanning mutagenesis of the aforementioned residues and assessed for the nuclease activity. Towards this, we designed a 100 bp target substrate harbouring a TTC PAM and performed the assay in the presence of Cascade/I-C and Cas3/I-C (Figure 2E-F). As compared to the wild type, mutants harbouring K743A, Q745A, and Y747A were found to be inactive (Figure 2F). The effect of these CTD mutations on CRISPR interference *in vivo* was tested in *E. coli* IC-1. This strain was transformed with pCas3/I-C harbouring the desired mutation in the CTD and the transformation efficiency was measured against pUC19 harbouring the target sequence. We anticipated that the active Cas3/I-C would target the pUC19 resulting in reduced transformation efficiency whereas the inactive Cas3/I-C would result in target evasion leading to high transformation efficiency. We observed that K743A, Q745A, and Y747A in CTD produced high transformants compared to WT, suggesting that these mutations indeed render the CRISPR interference inactive (Figure 2G & Supplementary Figure S7A).

### Resection of short double-stranded linear DNA by Cas3/I-C necessitates the presence of Cascade/I-C

Piqued by the observation that Cas3/I-C lacks intrinsic target specificity *in vitro* and retains one *in vivo*, we set out to resolve this apparent paradox. We hypothesised that the absence of “intrinsic” target specificity could be compensated by a mechanism that introduces “induced” target specificity. To test our hypothesis, we designed a 100 bp DNA substrate with “TTC” PAM at the 5’ end (Figure 2E). When Cas3/I-C alone was introduced, no apparent cleavage was noted (Figure 3A). This was surprising; however, it was consistent with our earlier observation (Figure 1D) that nuclease activity is attenuated for short DNA constructs (<400 bp). Led by the requirement of Cascade/I-C for CRISPR interference *in vivo*, we introduced the Cascade/I-C into the reaction. Remarkably, the target DNA – which was initially refractory to nuclease activity of Cas3/I-C – was cleaved in the presence of Cascade/I-C (Figure 3A and Supplementary Figure S5E). Intrigued by this, we asked what was bestowed by the Cascade/I-C on Cas3/I-C to target the DNA? Based on previous reports (21,32,33), we interpreted that the Cascade/I-C binding to target DNA ensues the generation of a DNA-RNA hybrid and a single-stranded DNA referred to as R-loop. Since Cas3/I-C requires 3’-overhang for loading (Figure 1F, Supplementary figure S2A, S2E-F), we hypothesised that the duplex nature of the target DNA perhaps abrogates the DNA binding. Therefore, it is likely that the ssDNA that was formed as a consequence of R-loop could then provide a platform for Cas3/I-C loading (Figure 3B). To test this, we designed a 100 bp bubble DNA construct that has no base pairing in the central region and that mimics the R-loop in the absence of Cascade/I-C (Figure 3C). Remarkably, we found that Cas3/I-C sliced the bubble DNA even in the absence of Cascade/I-C, suggesting that it is indeed the single-stranded region that is becoming the loading point for Cas3/I-C (Figure 3D and Supplementary Figure S5F).

**Figure 3.**
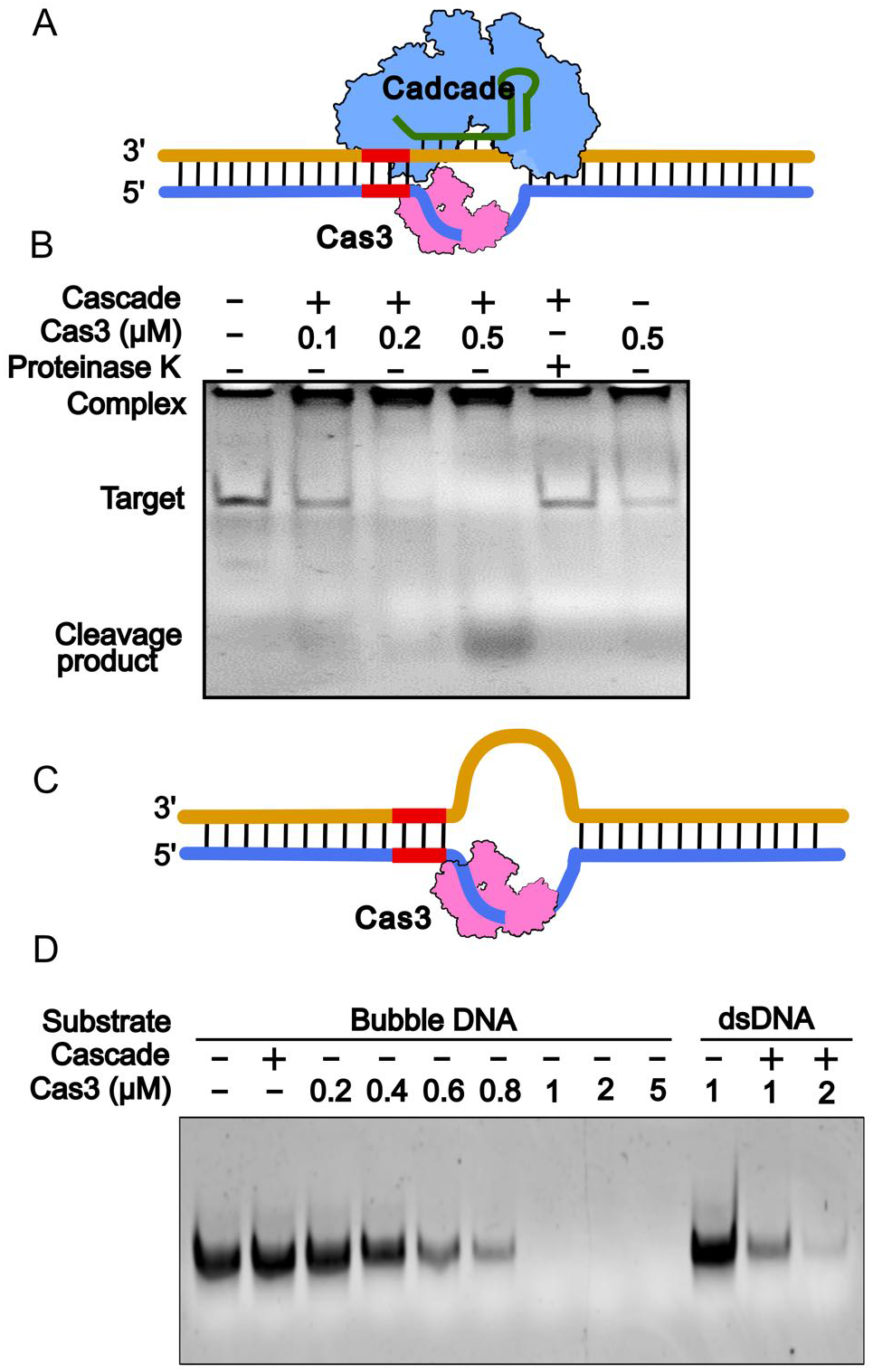
Cascade/I-C provides single-stranded DNA loop for Cas3/I-C binding. (A) A schematic model that suggests that Cascade/I-C binding to target DNA provides an ssDNA platform for Cas3 binding. The location of PAM (TTC in red) in the 100 bp construct was indicated. (B) DNA cleavage was tested on 20 % native PAGE with increasing concentration of Cas3/I-C in the presence of Cascade (lanes 2-4). Proteinase K was added to release bound DNA (lane 5). No cleavage was noticed in the absence of Cascade (lane 6). (C) A 100 bp bubble DNA construct that lacks base pairing in the centre was designed with PAM (TTC in red). The single-stranded DNA loop is suggested to facilitate Cas3/I-C binding in the absence of Cascade/I-C. (D) Bubble DNA was cleaved in the absence of Cascade/I-C (lanes 3-9) whereas dsDNA was cleaved only in the presence of Cascade/I-C (lanes 11-12) as visualised using 20% native PAGE.

### Cas3/I-C cleaves target via a reeling mechanism

The interaction between Cascade/I-C and Cas3/I-C facilitates the target DNA cleavage. After recruitment by Cascade/I-C, the helicase activity of Cas3/I-C may allow it to translocate on the ssDNA, unwinding the target DNA in 3’ to 5’ direction followed by cleavage via nuclease domain, eventually leading to the dissociation from Cascade/I-C. An alternative possibility is to remain associated with Cascade/I-C and reel in the target DNA, which may result in the formation of a loop in the target strand. In order to test, how Cas3/I-C unwinds double-stranded target DNA upon recruitment by Cascade/I-C, we designed a 100 bp dsDNA with a fluorophore (6-FAM) and a quencher (Iowa Black FQ). Non-target strand (NTS) was tagged with biotin at 5’ end and 6-FAM at 5^th^ nucleotide from the 5’ end. On target strand (TS), Iowa Black FQ (IB) was introduced proximal to the PAM at 29^th^ nucleotide from 3’ end (Figure 4A). Biotin was introduced at 5’ end with the presumption that Cas3/I-C would halt on encountering the biotin-streptavidin block thus sustaining 6-FAM quenching by Iowa Black FQ for a sufficiently long duration for measurements. Since 6-FAM and IB were far apart, the fluorescence of 6-FAM was not quenched. However, if Cas3/I-C reeled in the DNA, 6-FAM would come in proximity with IB, which in turn would quench the fluorescence. Target DNA was saturated with streptavidin and Cascade/I-C before adding Cas3/I-C and fluorescence intensity was measured to estimate the extent of quenching. Interestingly, we observed significant fluorescence quenching with increasing Cas3/I-C concentrations (0-5 μM) when Cascade/I-C was present. On the other hand, quenching was not observed when the reaction did not contain Cascade/I-C (dsDNA) or only non-target strand (ssDNA) was present (Figure 4B and supplementary figure S6F). Moreover, fluorescence quenching was observed only when ATP was supplied to Cas3/I-C in addition to Cascade/I-C (Figure 4C). Taken together, our result suggests that Cas3/I-C remain bound to Cascade/I-C and pulls in 6-FAM towards IB, which leads to fluorescence quenching.

**Figure 4.**
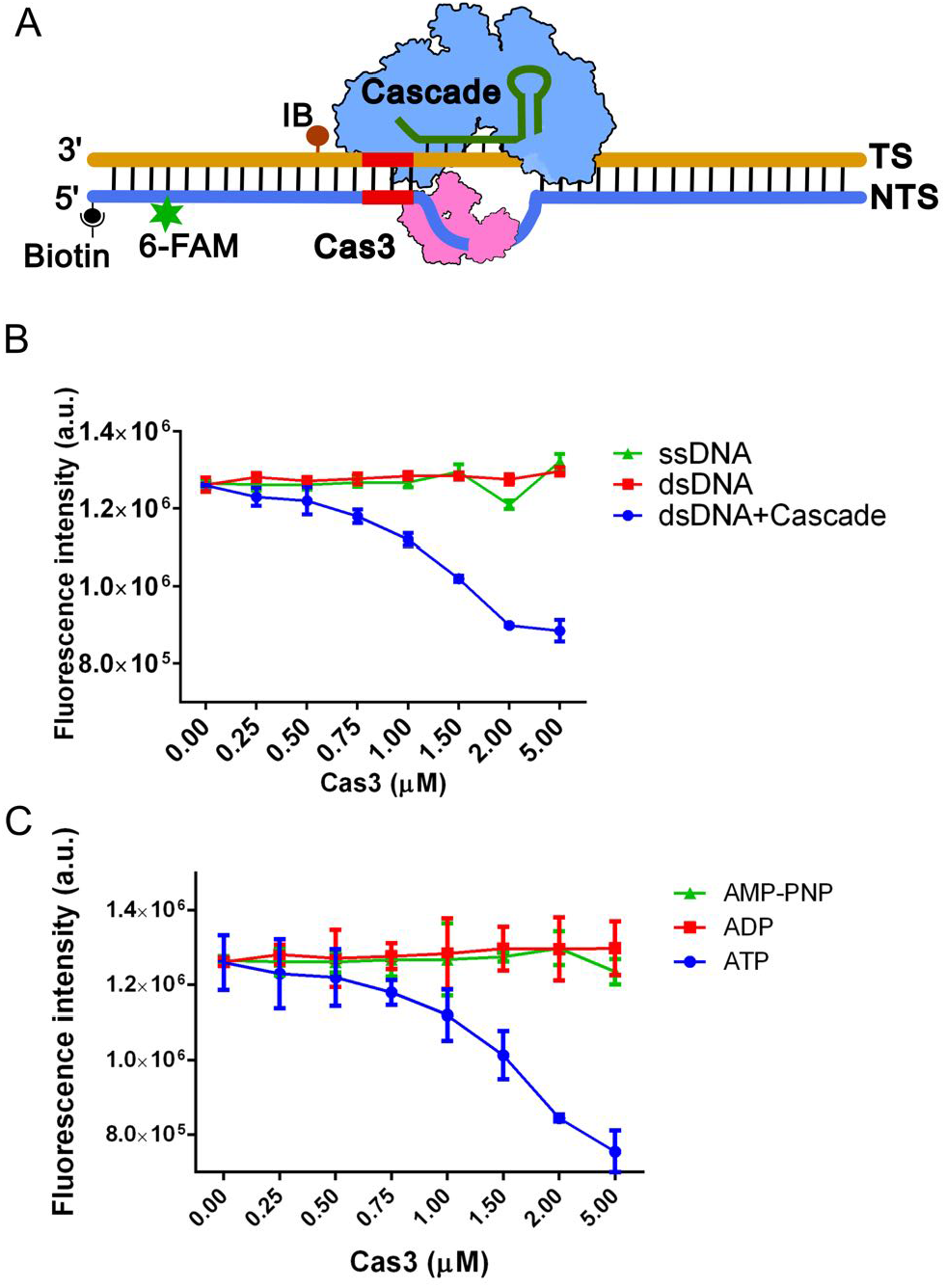
Cascade and Cas3 form a stable complex during interference. (A) Schematic representation of 100 bp DNA substrates with biotin (black dot) at 5’ end of the nontarget strand (NTS), 6-FAM (fluorophore, green star) at 5^th^ nucleotide, Iowa Black^®^ FQ (quencher, brown dot) at 28^th^ nucleotide on target strand (TS) and PAM sequence TTC (depicted in red colour). (B) Substrate mentioned above was incubated with or without Cascade/I-C (1 μM) and increasing concentration of Cas3/I-C (0-5 μM). A significant decline in fluorescence intensity is evident when both Cascade/I-C and Cas3/I-C were present. There was no apparent quenching when dsDNA and ssDNA were used in the absence of Cascade/I-C (C) Fluorescence quenching was observed in the presence of ATP but not when ADP and AMP-PNP were used.

### Stalling the Helicase motor of Cas3/I-C stimulates the Nuclease activity

Bacterial cytoplasm holds a repertoire of DNA binding proteins and therefore, Cas3/I-C during its processive nuclease-helicase activity is expected to encounter proteins such as single-stranded binding protein (SSB), polymerases, etc. In order to decipher what would happen if Cas3/I-C encounters such roadblocks, we intentionally introduced a stalling site in the target DNA. The stalling site comprises a biotin labelled nucleotide that binds to Streptavidin, which is expected to block the movement of Cas3/I-C helicase motor. We labelled the 5’ end of the non-target strand (NTS) with FAM and 12^th^ nucleotide from 5’ end with biotin (Figure 5A). Prior to the addition of Cas3/I-C, target DNA was incubated with Streptavidin and Cascade/I-C. Cas3/I-C was introduced with ATP/ADP/AMP-PNP and cleavage was monitored using denaturing PAGE (Figure 5B). We could discern the accumulation of a prominent ~55-60 nt band in absence of ATP. Since helicase motor is inactive in absence of ATP, we presume it to be a single-stranded nick generated by Cas3/I-C nuclease upon binding to NTS. Apart from the above observation, the presence of cleavage product (~55-60 nt) in the absence of nucleotide suggests that Cas3/I-C induced initial nick is independent of the nucleotide-bound state. Interestingly, in the presence of ATP, we observed a prominent band at ~40 nt; however, the band corresponding to ~55-60 nt was not perceptible. On the contrary, ~40 nt band was not visible when the biotin-streptavidin block was absent, even in the presence of ATP; however, we could spot the ~55-60 nt band (Figure 5B). This suggests that Cas3/I-C introduces a nick in the single-stranded region of NTS irrespective of the nucleotide-bound state.

**Figure 5.**
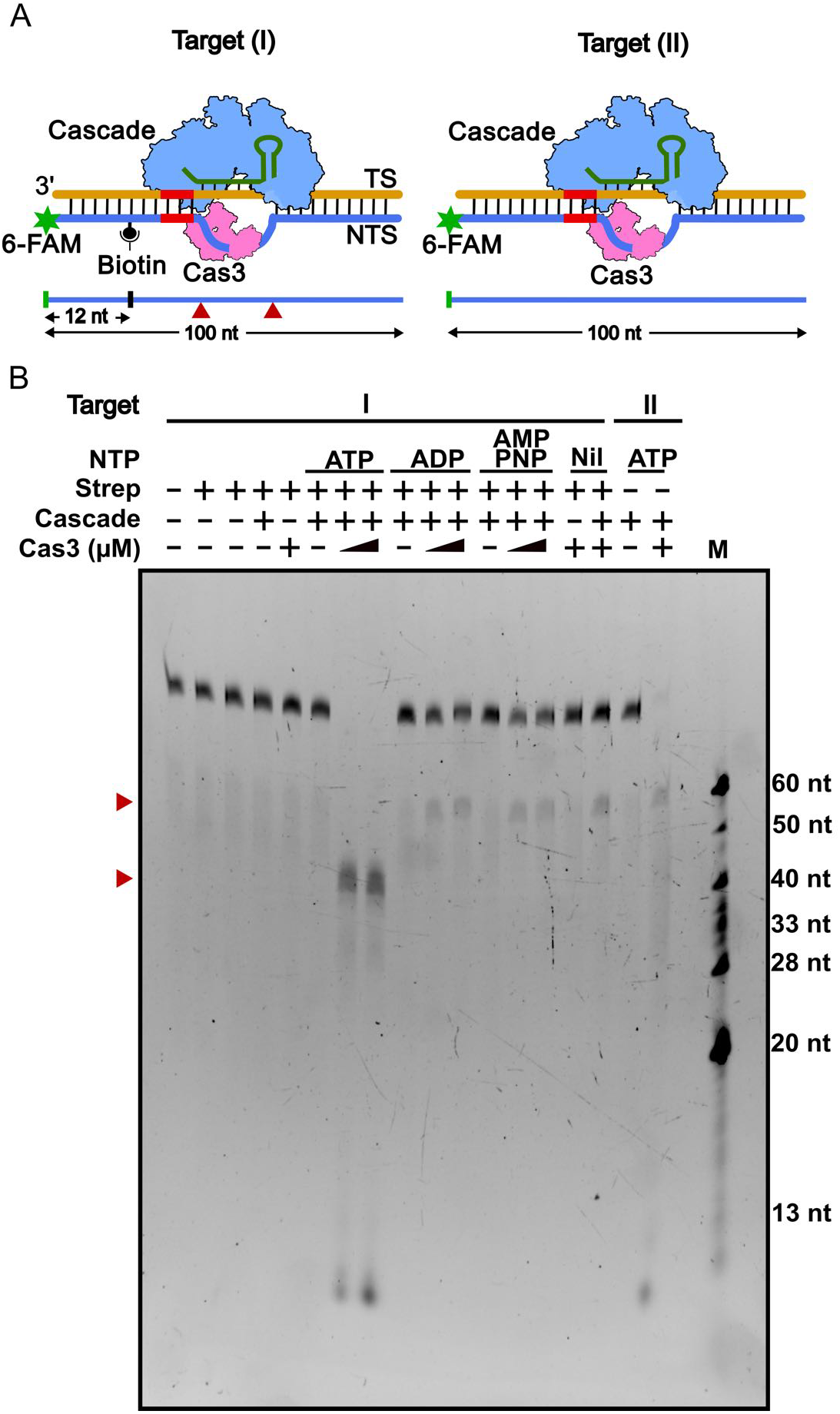
Stalling the translocation of helicase motor stimulates cleavage. (A) Schematic representation of target DNA (100 bp) with PAM (TTC in red) end-labelled with 6-FAM at 5’ end of the non-target strand and biotinylated at 12^th^ nucleotide (Target I). A similar target DNA without biotin (Target II) is also represented. Indicated in red arrow are the prominent cleavage sites. (B) Target (I) & (II) were incubated with streptavidin and Cascade/I-C to form interference complex. Cleavage was initiated by addition of Cas3 (0.2 and 0.5 μM) and 1 mM ATP/ADP/AMP-PNP. Prominent cleavages of the target (I) and (II) are indicated by a red arrow (approx. 60 nt and 40 nt). A 20% denaturing PAGE was used to assess the cleavage.

To account for the appearance of ~40 nt band specifically in the presence of ATP and biotin-streptavidin block, we hypothesized that the helicase motor of Cas3/I-C, in the presence of Cascade/I-C and ATP, would have reeled in the NTS from the loading point and a biotin-streptavidin roadblock could have triggered cleavage by the trailing nuclease domain. To test our hypothesis, we intentionally introduced similar stalling site in singlestranded DNA labelled with 6-FAM at 5’-end. Two constructs were made in which the biotin was labelled at 12^th^ (Target A) and 20^th^ (Target B) nucleotide positions, respectively (Figure 6A and 6D). We measured the change in fluorescence anisotropy of above-mentioned substrates upon addition of Cas3/I-C (Figure 6B, 6E and Supplementary figure S6C and S6D). Initially, we observed a slight increase in anisotropy for biotin-labelled DNA, suggesting early binding of Cas3/I-C to ssDNA. With time, we could discern a significant decline in anisotropy value in the absence or when ATP/ADP/AMP-PNP was present; however, the decay was steep when ATP was present. Remarkably, in the absence of biotin the anisotropy decay is not steep (Figure 6B & 6E). This decrease in anisotropy value with time suggests that DNA is fragmented possibly due to Cas3/I-C induced cleavage. In order to understand such cleavage, Cas3/I-C was mixed with ATP/ADP/AMP-PNP and bands were analysed using denaturing PAGE (Figure 6C and 6F). In line with above observations, two prominent bands at the top corresponding to ~30 and 40 nt for Target A (Figure 6C) and ~45 and 55 nt for Target B (Figure 6F), respectively, were perceptible in presence of ATP/ADP/AMP-PNP. For Target A, we posit that the ~30 nt band could arise by cleavage due to biotin block and the other ~40 nt could emerge by cleavage when a second Cas3/I-C gets stalled by the preceding Cas3/I-C. A similar scenario can be envisaged for Target B as well. Whereas in the absence of a biotin-streptavidin block, we could not observe any cleavage (Supplementary figure S6E). Interestingly, we also observe small sized bands (<10 nt), which suggests that few Cas3/I-C could trespass the biotin-streptavidin block and possibly get stalled by 6-FAM at the 5’ end (Figure 6C & 6F). Altogether the data suggest that when the helicase motor of Cas3/I-C encounters roadblock en route, the nuclease activity associated with HD domain is stimulated.

**Figure 6.**
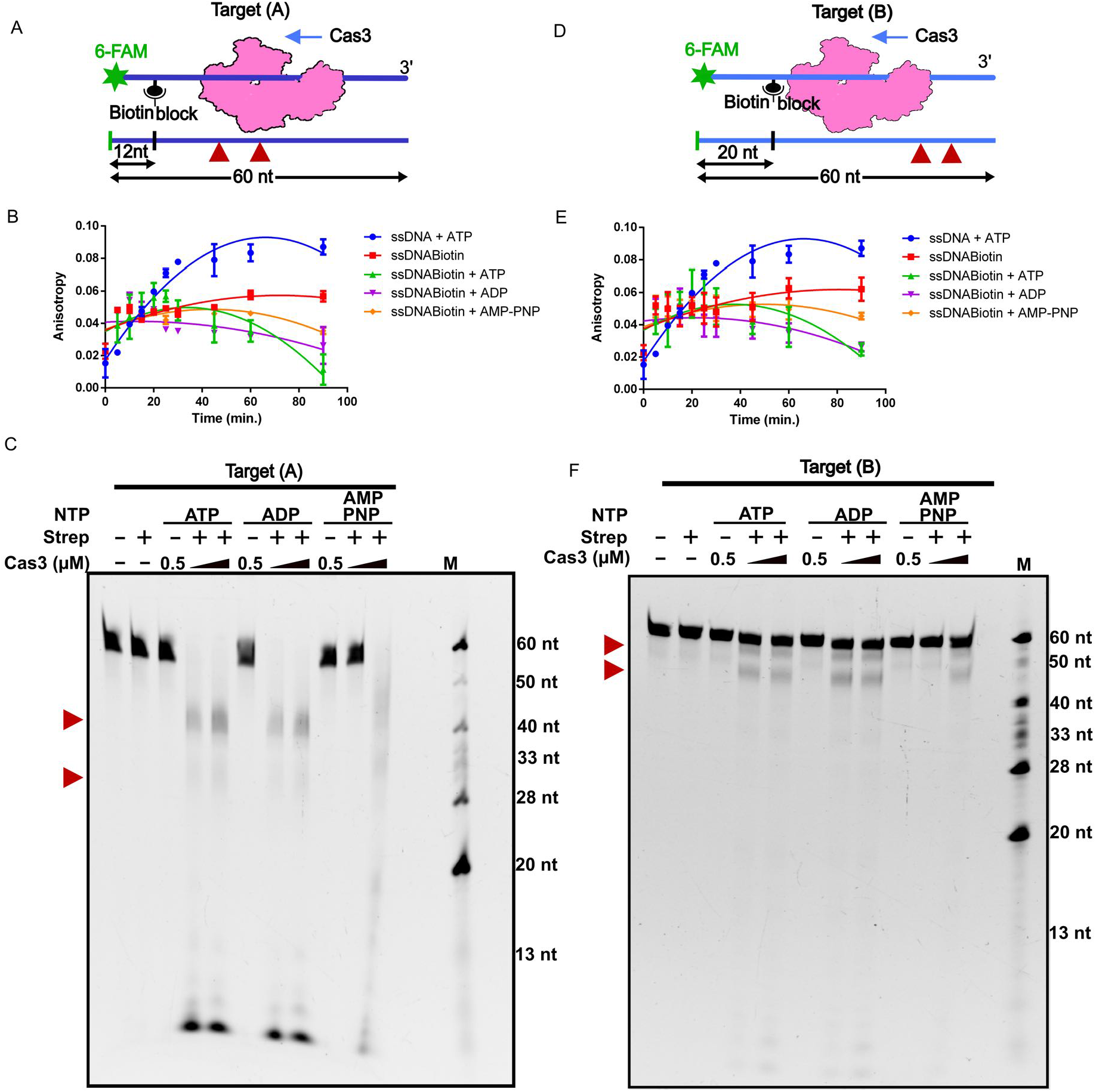
Roadblock in the translocation of Cas3/I-C stimulates cleavage. (A) & (D) A schematic representation of 60 nt 5’ 6-FAM labelled ssDNA with biotin at 12^th^ (Target A) and 20th nucleotide (Target B), respectively. (B) & (E) ssDNA mentioned above was incubated with 200 nM of Cas3/I-C for several time points and anisotropy measurements were recorded. With time, the decrease in anisotropy values was observed for ssDNA with a biotin roadblock. (C) & (F) ssDNA was pre-incubated with streptavidin before the addition of Cas3/I-C. Prominent DNA cleavage products were observed in the presence of ATP at ~40 nt in target A and ~50 nt in target B, indicated with a red arrow. In the presence of AMP-PNP higher Cas3 concentration was required for the cleavage. A 20% denaturing PAGE was used to study the cleavage pattern.

### Evolutionary conservation of target recognition and cleavage mechanism between type I-C and type I-E

Buoyed by the fact that it is the generation of single-stranded DNA as a by-product of R-loop that provides a platform for Cas3/I-C loading and target cleavage, we hypothesised that such mechanism could be pervasive among type I system irrespective of the differences in composition and architecture of Cascade. To test this, we chose Cascade/I-E – which is made up of five Cas subunits – as against Cascade/I-C, which comprises of three subunits only. Despite these differences, both Cascades facilitate the formation of R-loop and therefore, we anticipated that in such scenario it is possible to complement the Cascade/I-E with Cas3/I-C in order to promote the target cleavage *in vivo*. To test this, we created an *E. coli* strain (IC-2, Supplementary table S2) that harbours a spacer that targets pRSF-11 (T2, Supplementary table S3) and Cascade/I-E. In IC-2, the innate Cas3/I-E was deleted and therefore the intrinsic CRISPR/I-E machinery can recognise the target but it is deficient in target cleavage (Figure 7A). When IC-2 was challenged with T2, more number of transformants was observed (Figure 7B), suggesting that target cleavage is impaired in the absence of Cas3/I-E. However, when this strain was complemented with pCas3/I-C, there was a drastic reduction in the number of transformants, suggesting that T2 is efficiently targeted in the presence of Cas3/I-C (Figure 7B & Supplementary Figure S7B). To probe this further, the HD and DEXD/H domains of Cas3/I-C were co-expressed as independent domains for complementation *in vivo*. Remarkably, this strain showed less transformation efficiency that is comparable to WT, suggesting that these domains interact and produce functional Cas3/I-C *in vivo* during CRISPR interference. It is to be noted that HD and DEXD/H domains, when expressed separately, failed to restore CRISPR interference and it was only when they were co-expressed, T2 was targeted (Figure 7B). In line with this, pCas3/I-C, harbouring point mutation in lieu of catalytic residue in HD or Helicase domain is found to be inactive (Figure 7B). This suggests that despite the difference in the architecture of Cascade/I-C and Cascade/I-E, the Cas3 mediated target cleavage mechanism seems to be conserved between type I-C and type I-E.

**Figure 7.**
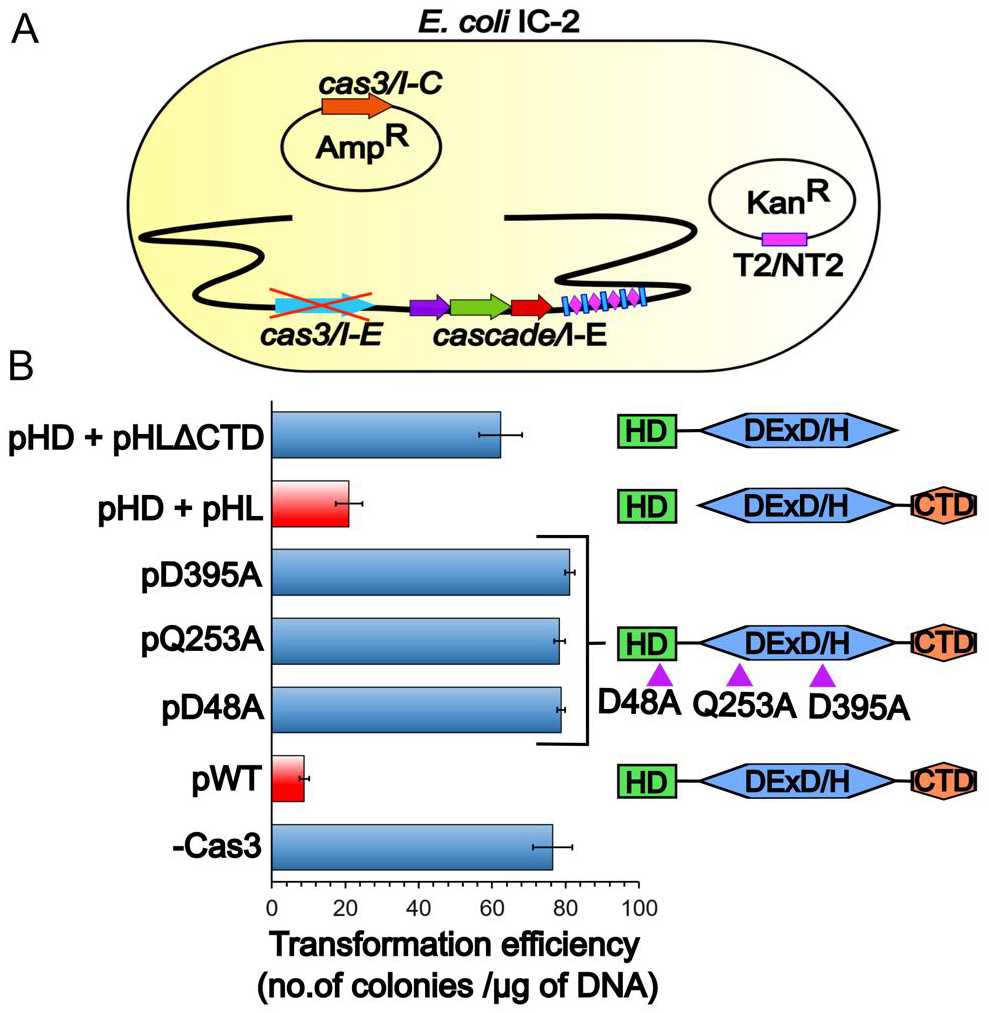
Cas3/I-C supplants Cas3/I-E in type I-E CRISPR-Cas system. (A) *E. coli* K-12 harbouring Cas3/I-E null mutant (IC-2) was used. IC-2 harbours spacer sequence targeting plasmid pRSF-11 (pT2). Cas3/I-C was heterologously expressed through IPTG inducible vector having ampicillin selection marker. Transformation efficiency was calculated after induction with 20 μM IPTG and 12 hr incubation. (B) Cas3/I-C and its variants were tested for interference activity using *E. coli* IC-2. Heterologous expression of Cas3/I-C (pWT) showed reduced transformation efficiency (indicated as a red bar). When the HD (pHD) and helicase (pHL) domains of Cas3/I-C were co-expressed as two separate constructs, the transformation efficiency mirrored that of wild type (indicated as a red bar). However, HD nuclease mutant (pD48A), helicase mutants (pQ253A and pD395A) and co-expressed constructs of HD and helicase without CTD (pHD + pHLΔCTD) showed high transformation efficiency. Error bar represents standard deviation measured from three independent trials. Triangles on the cylinders represent the location of the point mutations. The boundary of the domain variants are as follows: HD domain (pHD) 1-248; DExD/H domain (pHL) 249-800; pHD+pHLΔCTD 1-709.

## DISCUSSION

During CRISPR interference, precise recognition of target forms the cornerstone of self versus non-self-recognition. In Class II, a multi-domain single protein, *i.e*., Cas9 in type II-A, executes both target recognition and annihilation. However, in Class I/type I, Cascade recognises the target, whereas Cas3 cleaves the target. Unlike Class II, this division of labour between Cascade and Cas3 in Class I/type I suggests the possibility of additional regulatory control to the CRISPR based genome defence and therefore alludes to the prevalence of type I among the prokaryotes. In this context, one could perceive that Cascade and Cas3 should have “intrinsic” target specificity to obviate any collateral damage on the host. In the case of Cascade, this is satisfied by complementary base pairing between crRNA and the target DNA. In line with this, our data suggest that the differences in the architecture and composition of Cascade among type I are unlikely to wield high influence in determining the specific recognition of the target DNA – a role that is exquisitely delegated to crRNA (Figure 7). On the contrary, we noted that the nuclease activity of Cas3 in type I is highly promiscuous (Figure 1) – which is counter-intuitive – and this has an intrinsic potential for indiscriminate host-targeting (43,44,64). Intriguingly, in type I variants such as type I-C and type I-E, though the composition and architecture of Cascade seem to differ, the overall architecture of Cas3 is broadly conserved, suggesting that Cas3 recognizes the target DNA via a “generic” mechanism and it is likely that this is independent of the structure of Cascade. Piqued by this, we set out to unravel the underlying mechanism by which Cas3 selectively degrades the target DNA without inflicting collateral damage.

In Cas3/I-C, the requirement of HD, DEXD/H and CTD domains during CRISPR interference suggests that the concerted domain-domain crosstalk plays a pivotal role in regulating the nuclease activity (Figure 2, 7 & Supplementary Figure S7). Strikingly, both HD and DEXD/H domains require single-stranded DNA substrates, which are transiently formed during highly orchestrated events such as replication and transcription. Therefore, Cas3 is unlikely to have access to its preferred single-stranded substrates most part of the time and this may preclude any inadvertent self-targeting. The absence of cleavage of small double-stranded DNA (< 400 bp) as against plasmid DNA lends support to this conjecture (Figure 1). Due to the typical *in vitro* setting by which plasmid DNA is isolated by employing denaturation and renaturation cycles as well as given its large size, it is likely that some regions retain single-stranded state even after renaturation. Moreover, pQE2 is found to be negatively supercoiled, which may also produce a single-stranded region. These singlestranded regions could become the substrate for Cas3 mediated nicking and its subsequent loading points. Whereas shorter DNA fragment is likely to have efficient renaturation and this can pre-empt Cas3 loading. Also, this scenario is likely to get reversed when the Cascade mediated recognition of the foreign DNA leads to the formation of R-loop and this sets the stage for Cas3 to access the single-stranded substrate. Our data suggest that the recruitment of Cas3 towards R-loop is not solely contingent on Cascade, which is in part driven by the single-stranded region (Figure 3C & 3D).

Nonetheless, the modelled structure of Cas3/I-C based on Cas3/I-E suggests that the active site of HD nuclease is oriented towards the helicase domain and therefore it is unlikely to access the ssDNA directly. Consequently, it appears that the ssDNA has to enter the HD active site via the helicase domain only. However, our data suggest that upon recruitment to the target site, Cas3/I-C nicks non-target strand at ~30-35 nt from PAM (Figure 5). This initial nick in the absence of ATP points towards the direct binding of nuclease domain at PAM distal region of R-loop, which then renders a loading point for the helicase domain (49). This also concurs with the interpretation from the recent Cryo-EM structure of Cascade/I-E and Cas3/I-E (46). Our data further suggest that though Cas3/I-C lacks DNA unwinding activity in the absence of ATP, it can translocate on the ssDNA (Figure 6). In the presence of ATP, it seems that Cas3/I-C unwinds the target DNA as well as channelises the displaced ssDNA towards the active site of the HD nuclease through the helicase motor activity. This implies that in the absence of the helicase motor activity, the DNA is unlikely to reach the HD nuclease. This is in line with the current understanding on Cas3/I-E mediated cleavage and explains the requirement of the helicase domain for CRISPR interference (Figure 2 & 5) (35,46). We envisage another possible scenario wherein despite the helicase motor and HD nuclease being active, mutations in the channel forming residues are likely to derail the conduit so that the displaced ssDNA is unlikely to be threaded towards the active site of the HD nuclease resulting in the loss of target cleavage. The R-loop has no free 3’end for helicase loading and therefore threading the ssDNA into the helicase domain creates a topological problem. Previous studies on Yeast DEAD Box Protein Mss116p demonstrates the modulation of helicase core activity by CTD (65,66). Similarly, we identified a few critical residues in CTD of Cas3/I-C – which appears to be unique with respect to type I-C – and that possibly stabilises the DNA-bound helicase core thus bolting the channel for DNA translocation (Figure 2 & Supplementary Figure S2E and S4).

Presumably, we posit that the incessant movement of DNA towards the HD nuclease would not provide enough residence time for the alignment of catalytic residues. This is likely to impede the nuclease activity as long as the helicase motor is active (Figure 4-5 & Supplementary Figure S4B and S6E). Therefore, whenever there is a blockade for helicase movement, it is likely that the catalytic residues of HD nuclease have sufficient residence time to be suitably oriented towards the scissile bond (Figure 4-5 & Supplementary Figure 4B). This appears to be an ingenious strategy to time the cleavage. Our data on Cas3/I-C are in agreement with the recent finding where stalling of Cas3/I-E translocation at roadblocks stimulates nuclease activity which is otherwise reported to be sparse (49,67). After the Cas3/I-C loading onto the single-stranded region, we envisage two modes of helicase motor activity: (i) translocation - Cas3/I-C is mobile with respect to the bound ssDNA and (ii) reeling - Cas3/I-C is stationary with respect to the bound ssDNA. However, in line with the previous report on Cas3/I-E (49), our data showed the cleavage pattern that concurs with a DNA reeling mechanism wherein Cas3/I-C has an intimate interaction with Cascade during DNA unwinding (Figure 4 & Supplementary Figure 6F). In this set up – where the mobility of the first Cas3/I-C molecule is locked due to interaction with Cascade/I-C – a caravan consisting of multiple Cas3/I-C that can load onto the R-loop is likely to collide with each other as well as with the one that interacts with Cascade/I-C to inflict multiple cleavages (Figure 6). While the rationale for adopting reeling over translocation is not abundantly clear, it is tempting to speculate that reeling allows Cas3/I-C to inflict cleavage that is proximal to the interference effector complex and endows with an apparent target site specificity (Figure 8).

**Figure 8.**
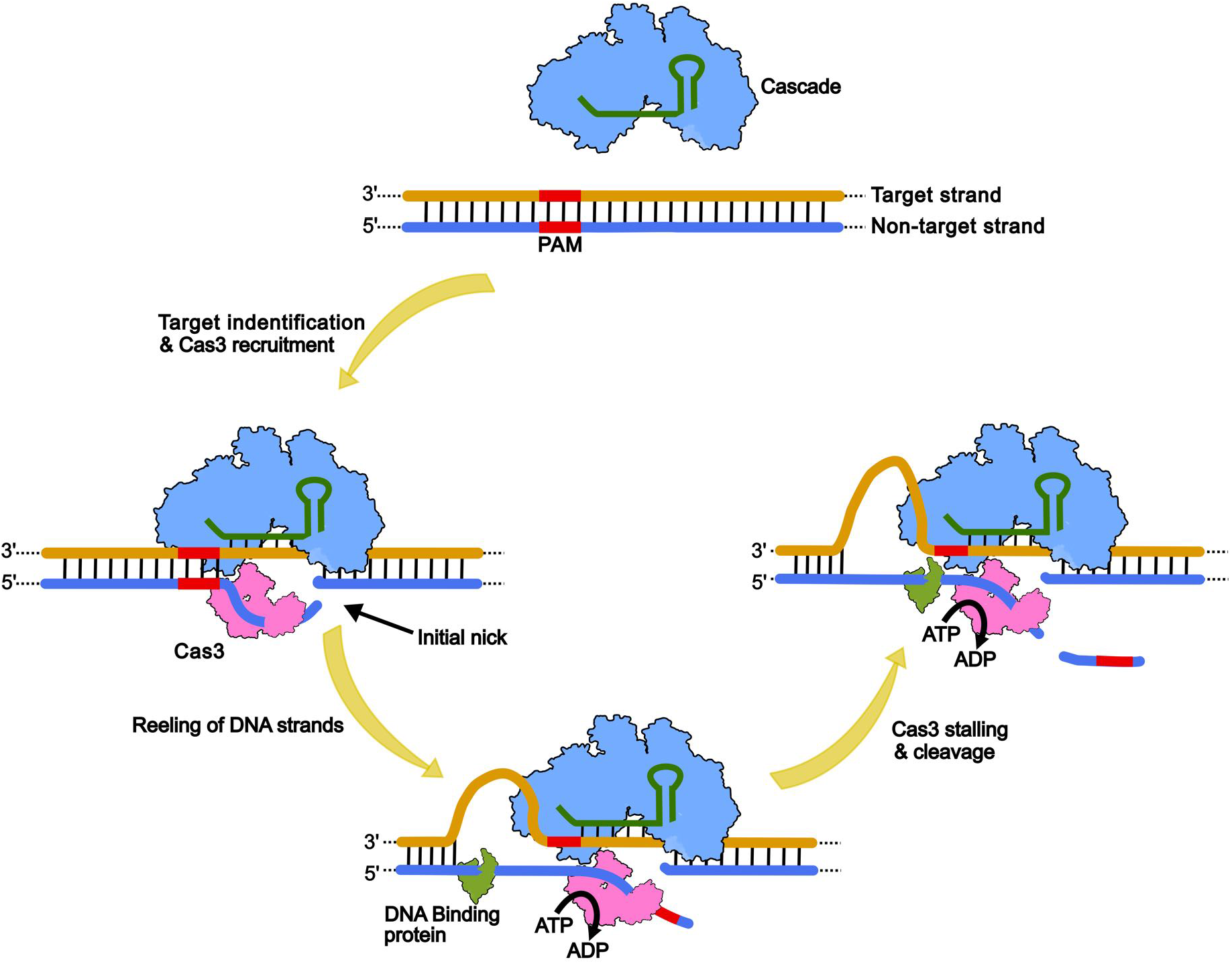
A schematic model representing the possible scenarios leading to Cas3 mediated target DNA cleavage. Cascade through base pairing between crRNA and target DNA locates the invading foreign genetic element, thereby forming an R-loop. The ensuing conformational change in the large subunit of Cascade facilitates its binding to Cas3 followed by HD domain-mediated single-stranded nick on the non-target strand. This promotes the loading of the helicase domain of Cas3 onto the ssDNA of the non-target strand. Based on the longevity of Cascade-Cas3 interaction, we propose that Cas3/I-C remains attached to the Cascade as a stable complex and reels DNA (DNA looping on target strand) in, utilising energy from ATP. Subsequently, Cas3 may hit a roadblock (e.g. DNA binding proteins), which can trigger further DNA cleavage. Here, the cleavage site is likely to be in proximity to the Cascade.

The fact that Cas3/I-C –albeit exhibiting differences such as the presence of highly conserved residues in CTD – shows functional similarities with respect to Cas3/I-E has raised a question how this functional equivalence could exist in harmony with Cascade/I-C –which exhibits compositional and architectural differences with Cascade/I-E. Strikingly, the complementation of Cas3/I-C in lieu of Cas3/I-E in type I-E system *in vivo* shows that the target recognition and cleavage mechanism is conserved between type I-C and I-E (Figure 7). When we compared the modelled structure of Cas3/I-C from *B. halodurans* (BhaCas3/I-C) (Figure 2D and Supplementary figure S3C(i)) and Cas3/I-E from *E. coli* (EcoCas3/I-E) based on the recent cryo-EM structure of Cas3/I-E from *Thermobifida fusca* (TfuCas3/I-E) (35), we could discern that Cas3/I-C shows significant sequence and structural similarities with Cas3/I-E from *E. coli* (EcoCas3/I-E), if not complete, in interfaces I to IV (Supplementary figure S3C and S8). Further, we attribute the reported incompatibility for hetero-assembly between TfuCas3/I-E and EcoCascade/I-E (46) to the following factors: (i) the reconstitution of hetero-assembly was attempted *in vitro*. Therefore, it is quite likely that the lack of cellular milieu could be a possible deterrent for hetero-assembly. (ii) TfuCas3/I-E and EcoCascade/IE are not thermo-compatible. The former is thermophilic and the latter is mesophilic. They have undergone different adaptation attuned with the thermal conditions. (iii) In our study, type I-C and I-E are not co-existing as composite CRISPR-Cas system within the same organism. This precludes any selection pressure to avert cross-over Cascade-Cas3 interaction between type I-C and I-E. Thus, our data suggest that despite the compositional and architectural differences seen between Cascade/I-C and Cascade/I-E, Cas3 recruitment and target cleavage mechanism seem to be conserved, at least in I-C and I-E systems, if not all type I.

## CONCLUDING REMARKS

It is quite remarkable to note that Cas3 – albeit being a promiscuous nuclease – is induced towards achieving its target specificity by a series of tightly coordinated events that segregate its nuclease and helicase activities in space and time. Instead of producing a new effector nuclease with intrinsic target specificity, the type I CRISPR-Cas system has adopted an existing generic helicase-cum-nuclease module and further evolved ways and means to nudge its unrestrained activity towards specifically targeting the foreign DNA. Evidently, this molecular metamorphosis of Cas3 is turning out to be a paradigm of resource optimisation for CRISPR immunity.

## SUPPLEMENTARY DATA

Supplementary Data are available online.

## ACKNOWLEDGEMENT

Vector pOSIP-CT (Addgene #45981) was a kind gift from Drew Endy and Dr Keith Shearwin; pST-KT was gifted by Dr Vinay K. Nandicoori; p1R (Addgene #29664), and p13SR (Addgene #48328) were provided by Scott Gradia; pKD46 (CGSC #7739) was a kind gift from Barry L. Wanner; *E. coli* IYB5101 was a gift from Udi Qimron. We acknowledge the geniality of the aforementioned scientists for sharing their plasmids and bacterial strains. We thank all members of the MAB lab for their critical comments and suggestions.

## FUNDING

This work was supported in part by grants from Department of Biotechnology (DBT) [BT/PR15925/NER/95/141/2015, BT/08/IYBA/2014/05, BT/406/NE/UEXCEL/2013, BT/PR5511/MED/29/631/2012 and BT/341/NE/TBP/2012] and Science and Engineering Research Board (SERB) [YSS/2014/000286].

## CONFLICT OF INTEREST

The authors declare no competing interests.

