## Supplementary Information for "Cas3 Mediated Target DNA Recognition and Cleavage is Independent of the Composition and Architecture of Cascade Surveillance Complex"

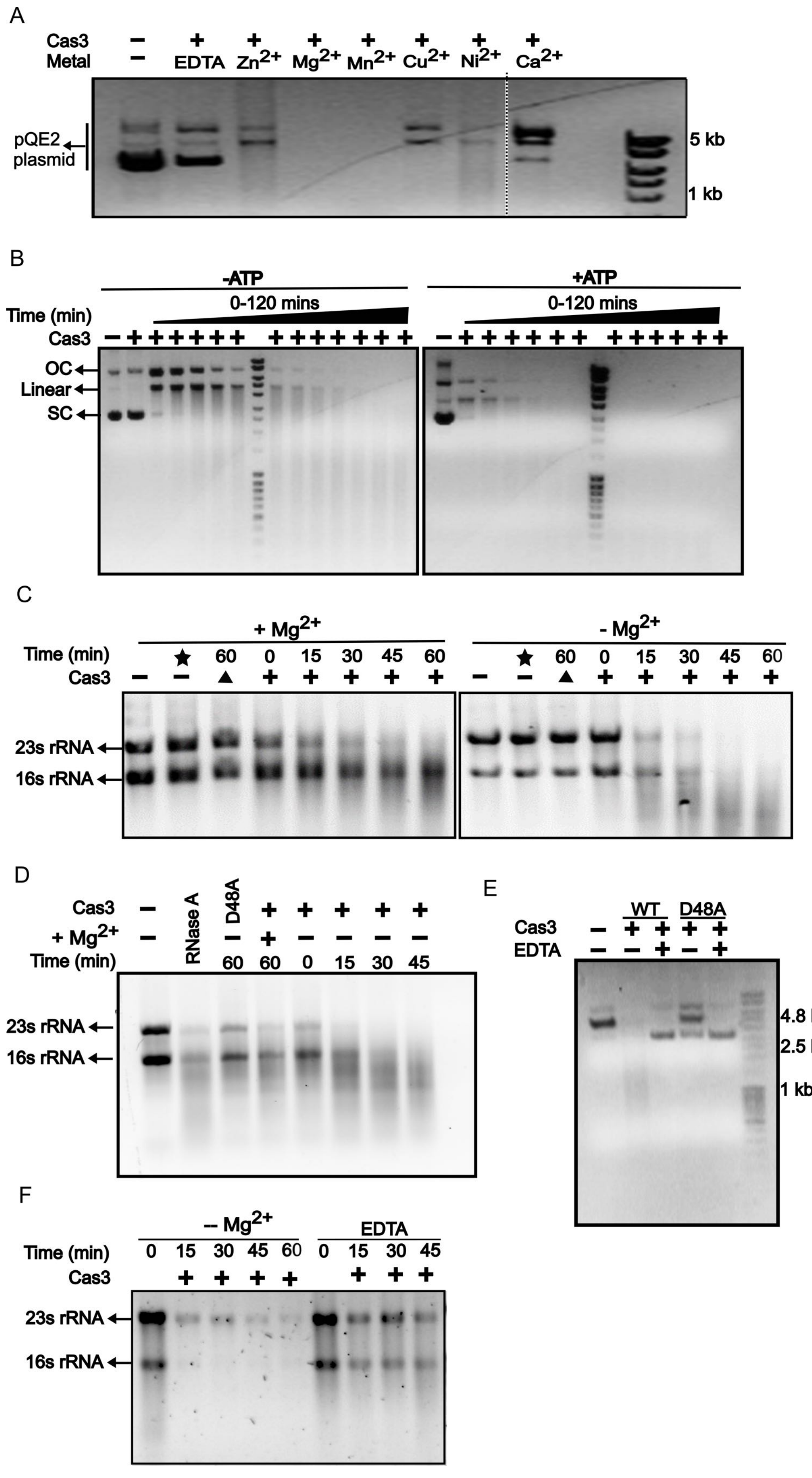
A
Cas3
Metal
EDTA
Zn2+
Mg2+
Mn2+
Cu2+
Ni2+
Ca2+
pQE2 plasmid
5 kb
1 kb
B
-ATP
+ATP
Time (min)
0-120 mins
Cas3
OC
Linear
SC
C
+ Mg2+
- Mg2+
Time (min)
60 0 15 30 45 60
Cas3
23s rRNA
16s rRNA
D
Cas3
+ Mg2+
Time (min)
RNase A
D48A
60 60 0 15 30 45
23s rRNA
16s rRNA
E
WT
D48A
Cas3
EDTA
4.8 kb
2.5 kb
1 kb
F
– Mg2+
EDTA
Time (min)
0 15 30 45 60 0 15 30 45
Cas3
23s rRNA
16s rRNA

**Figure S1: Nuclease activity of Cas3/I-C *in vitro*,** Related to Figure 1

(A) Cas3/I-C mediated DNA cleavage is dependent on the divalent metal ion. Addition of 500 nM Cas3/I-C in presence of $Mg^{2+}$ and $Mn^{2+}$ in the reaction mixture showed prominent cleavage whereas $Ni^{2+}$, $Zn^{2+}$, $Cu^{2+,}$ and $Ca^{2+}$ were less effective to a varying extent. EDTA inhibited the cleavage drastically. The dotted line indicates a discontinuity in the gel for the purpose of clarity.

(B) Time-dependent cleavage of plasmid DNA (pQE2) (0-120 mins) by Cas3/I-C shows that nuclease activity is enhanced in the presence of ATP, suggesting a functional overlap between the DEXD/H helicase and HD nuclease domains.

(C) 16S and 23S rRNAs isolated from *B. halodurans* were used as substrates for Cas3/I-C in the presence and absence of $Mg^{2+}$ and the reaction was monitored for several time points (0-60 mins). The absence of $Mg^{2+}$ made rRNAs more susceptible to cleavage. Since $Mg^{2+}$ is required for RNA folding, this enhanced cleavage is presumably due to the destabilization of RNA structure in the absence of $Mg^{2+}$. 'Triangle' denotes Cas3/I-C, which was rendered nuclease inactive after heat treatment at 80 ºC for 20 mins. Star symbol signifies that rRNA was incubated in the reaction buffer for 60 mins without the addition of Cas3/I-C to negate the possibility of nuclease contamination.

(D) 16S and 23S rRNAs were used as a substrate for Cas3/I-C WT and Cas3/I-C nuclease mutant (D48A). We have incubated rRNA substrate for various time points. Nuclease activity was abrogated by the nuclease domain mutation D48A. rRNA was incubated with RNase A for 60 mins.

(E) Plasmid DNA (pQE2) was incubated with Cas3/I-C WT and nuclease domain point mutant (D48A). DNA cleavage was severely attenuated with Cas3/I-C D48A.

(F) To check metal-dependent cleavage of RNA, 16S and 23S rRNA were used as a substrate for Cas3/I-C in presence of 10 mM EDTA and cleavage was monitored for several time points. Presence of EDTA inhibited rRNA cleavage, which suggests that Cas3/I-C co-purifies with a bound metal ion.

Note: All the samples from supplementary figure S1A to S1F are visualized using 0.8% agarose gel electrophoresis stained with ethidium bromide.

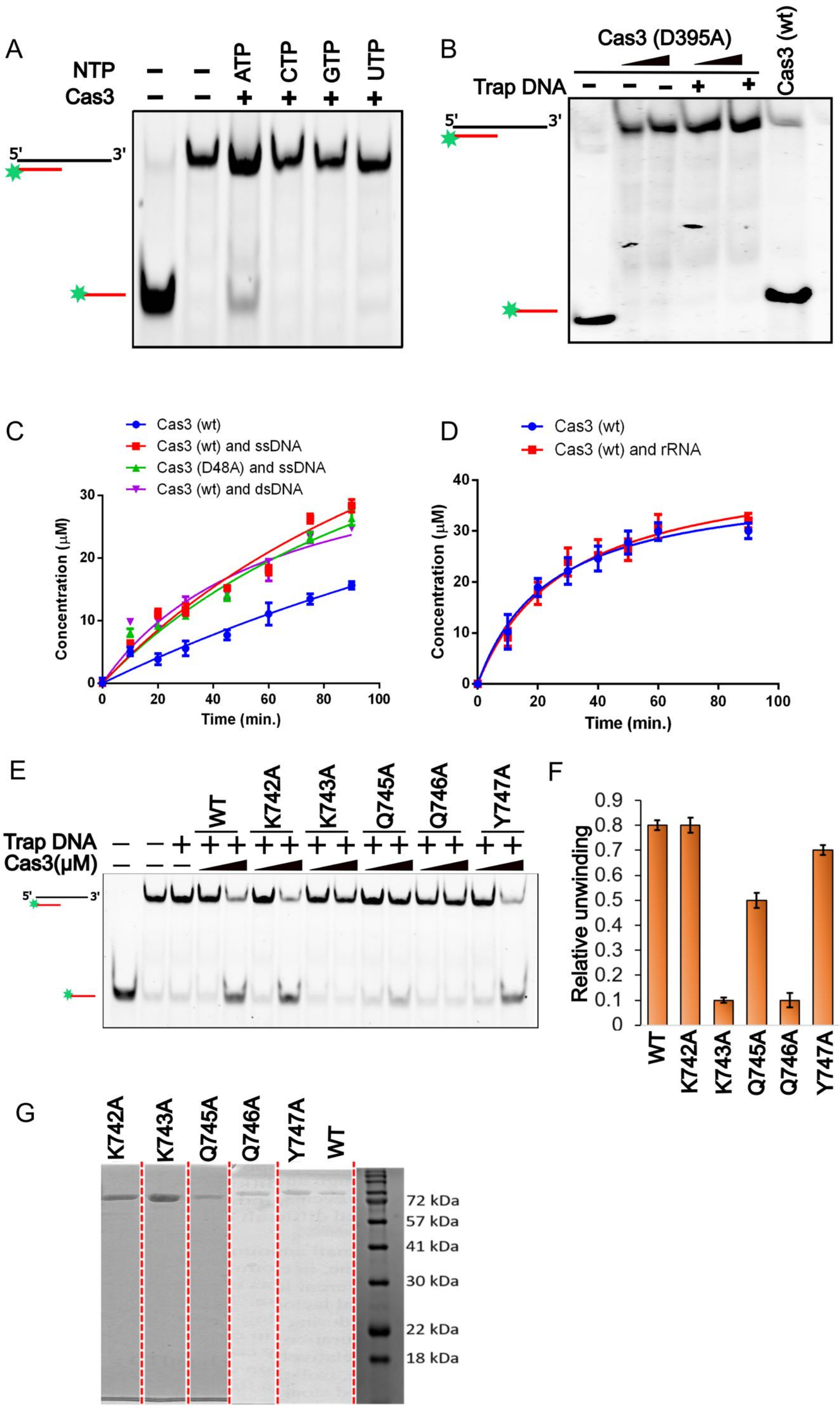


**Figure S2: Helicase activity of Cas3/I-C and its variants,** Related to Figure 1 and 2

(A) Cas3/I-C unwinds DNA utilizing ATP as the sole energy source. In the presence of other nucleotide triphosphates (CTP, GTP, and UTP), no unwinding was observed.

(B) Cas3/I-C helicase domain mutant (D395A) was unable to unwind partial duplex. We have used 250 nM and 500 nM Cas3/I-C concentration both with and without trap DNA.

(C) & (D) Rate of ATP hydrolysis is enhanced in the presence of nucleic acids. Cas3/I-C (wt) is shown to hydrolyse ATP in the absence of any nucleic acid substrate. Addition of M13mp18 (ssDNA) and 70 bp partial duplex DNA with 3' overhang (dsDNA) enhanced ATP hydrolysis significantly as seen in case of both Cas3/I-C (wt) and nuclease inactive Cas3/I-C (D48A). Whereas the rate of ATP hydrolysis remained unchanged when rRNAs (16S and 23S) were used as substrates with Cas3/I-C (wt). Error bar represents standard deviation measured from three independent trials.

(E) Fluorescently labelled partial duplex DNA (with 3' overhang) was used to test the helicase activity of CTD mutants. Trap DNA was provided in excess to avoid re-annealing of the displaced strand.

(F) Relative unwinding signifies the percentage of the released strand in 15 mins as observed in (E). DNA band intensities from (E) were quantified using Image Lab software (Bio-Rad). Error bar represents standard deviation calculated from two independent trials.

(G) Strep-tagged Cas3/I-C (93 kDa) was purified using multiple steps of purification. The figure shows the final preparation of Cas3/I-C and its variants employed in various assays in this study. The red dotted line indicates a discontinuity in gel shown for the purpose of clarity. Samples were run on 15% SDS-PAGE.

Note: Samples in supplementary figures S2A, S2B and S2E were visualized using 15 % native PAGE at 4 ºC.

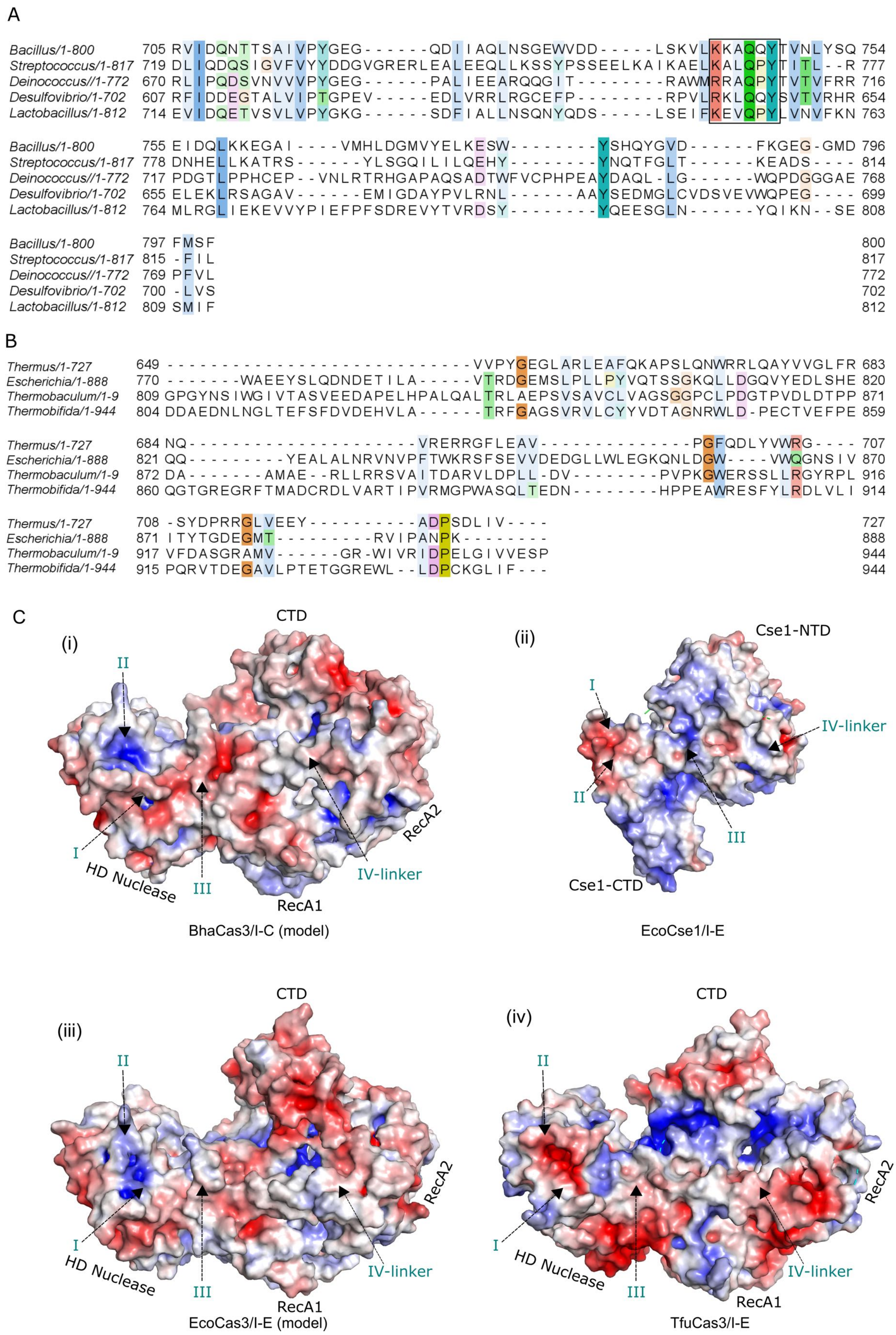
A
Bacillus/1-800 705 RVIDQNTTSAIVPYGEG------QDIIAQLNSGEWVDD-----LSKVLKKAQQYTVNLYSQ 754
Streptococcus/1-817 719 DLIQDQSIGVFVYYDDGVGRERLEALEEQLLKSSYPSSEELKAIKAELKALQPYTITL--R 777
Deinococcus//1-772 670 RLIPQDSVNVVVPYGEG------PALIEEARQQGIT--------RAWMRRAQPYTVTVFRR 716
Desulfovibrio/1-702 607 RFIDDEGTALVIPTGPEV-----EDLVRRLRGCEFP--------RPVLRKLQQYSVTVRHR 654
Lactobacillus/1-812 714 EVIDQETVSVLVPYGKG------SDFIALLNSQNYQDS-----LSEIFKEVQPYLVNVFKN 763
Bacillus/1-800 755 EIDQLKKEGAI----VMHLDGMVYELKESW--------YSHQYGVD------FKGEG-GMD 796
Streptococcus/1-817 778 DNHELLKATRS------YLSGQILILQEHY--------YNQTFGLT------KEADS---- 814
Deinococcus//1-772 717 PDGTLPPHCEP--VNLRTRHGAPAQSADTWFVCPHPEAYDAQL-LG------WQPDGGGAE 768
Desulfovibrio/1-702 655 ELEKLRSAGAV------EMIGDAYPVLRNL------AAYSEDMGLCVDSVEVWQPEG---- 699
Lactobacillus/1-812 764 MLRGLIEKEVVYPIEFPFSDREVYTVRDSY--------YQEESGLN------YQIKN--SE 808
Bacillus/1-800 797 FMSF 800
Streptococcus/1-817 815 -FIL 817
Deinococcus//1-772 769 PFVL 772
Desulfovibrio/1-702 700 -LVS 702
Lactobacillus/1-812 809 SMIF 812
B
Thermus/1-727 649 ----------------------------VVPYGEGLARLEAFQKAPSLQNWRRLQAYVVGLFR 683
Escherichia/1-888 770 -------WAEEYSLQDNDETILA-----VTRDGEMSLPLLPYVQTSSGKQLLDGQVYEDLSHE 820
Thermobaculum/1-9 809 GPGYNSIWGIVTASVEEDAPELHPALQALTRLAEPSVSAVCLVAGSGGPCLPDGTPVDLDTPP 871
Thermobifida/1-944 804 DDAEDNLNGLTEFSFDVDEHVLA------TRFGAGSVRVLCYYVDTAGNRWLD-PECTVEFPE 859
Thermus/1-727 684 NQ---------------------VRERRGFLEAV--------------PGFQDLYVWRG---- 707
Escherichia/1-888 821 QQ---------YEALALNRVNVPFTWKRSFSEVVDEDGLLWLEGKQNLDGW----VWQGNSIV 870
Thermobaculum/1-9 872 DA-------AMAE--RLLRRSVAITDARVLDPLL-DV---------PVPKGWERSSLLRGYRPL 916
Thermobifida/1-944 860 QGTGREGRFTMADCRDLVARTIPVRMGPWASQLTEDN---------HPPEAWRESFYLRRDLVLI 914
Thermus/1-727 708 -SYDPRRGLVEEY-----------ADPSDLIV---- 727
Escherichia/1-888 871 ITYTGDEGMT----------RVIPANPK-------- 888
Thermobaculum/1-9 917 VFDASGRAMV------GR-WIVRIDPELGIVVESP 944
Thermobifida/1-944 915 PQRVTDEGAVLPTETGGREWL--LDPCKGLIF--- 944
C
(i)
CTD
II
I
III
IV-linker
HD Nuclease
RecA1
RecA2
BhaCas3/I-C (model)
(ii)
Cse1-NTD
I
II
III
IV-linker
Cse1-CTD
EcoCse1/I-E
(iii)
CTD
II
I
III
IV-linker
HD Nuclease
RecA1
RecA2
EcoCas3/I-E (model)
(iv)
CTD
II
I
III
IV-linker
HD Nuclease
RecA1
RecA2
TfuCas3/I-E

**Figure S3: Comparison of Cascade interacting interface between Cas3/I-E and Cas3/I-C,** Related to figure 2 and figure 7

(A) Multiple sequence alignment corresponding to CTD of Cas3/I-C from select organisms is shown. The sequences were aligned using MUSCLE. The motif "KKAQQY" that shows high conservation among the orthologs is shown within the black box. The location of these residues in the homology model of Cas3/I-C suggests a possible role to stabilize the channel for target DNA. Residues in the alignment are highlighted based on the extent of conservation using Jalview (1).

(B) Cas3 sequences from type I-E systems were aligned using MUSLCE. In type I-E system, the motif "KKAQQY" is absent, suggesting some differences in the functionality of CTD between type I-C and I-E.

(C) Four interfaces on BhaCas3/I-C (i) and EcoCas3/I-E (iii) which were modelled using Cas3/I-E (PDB ID: 6C66), are shown in blue colour. These interfaces are identified as contact points between TfuCas3/I-E (iv) and Cse1 subunit of TfuCascade/I-E (2). The surface electrostatics of EcoCse1/I-E (ii) is shown along with expected Cas3 contact interfaces. Surface charges on BhaCas3/I-C (i) and EcoCas3/I-C (iii) shows similarities at interfaces I, II and III. Unlike TfuCas3/I-E (iv) and EcoCse1/I-E (iii), the charge compatibility between BhaCas3/I-C (i) and EcoCse1/I-E (iii) (mainly interfaces II and III) explains why there is a hetero-assembly of BhaCas3/I-C and EcoCascade/I-E *in vivo* in our study.

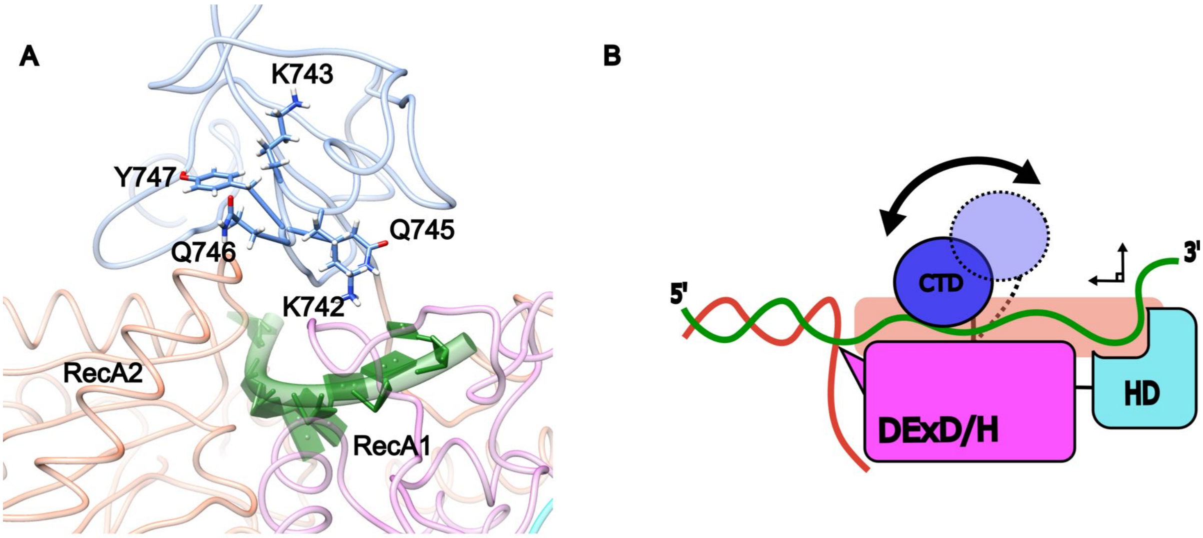


**Figure S4: Homology model of Cas3/I-C CTD based on Cas3/I-E structure [PDB ID: 4QQX]** Related to figure 2 and 3

(A) Cas3/I-C was modelled using Cas3/I-E [PDB ID: 4QQX] as a template using I-TASSER. Residues selected for mutations in this study are indicated in the figure. The position of these mutations suggests their role in locking CTD after Cas3/I-C is bound to target DNA during interference.

(B) The modelled structure of Cas3/I-C as well as available Cas/I-E structures suggests that the uncharacterized CTD rests on top of DExD/H domain, providing a channel for ssDNA to enter, thereby stabilizing the interaction between DNA and Cas3/I-C. CTD (shown in blue) is linked to the helicase domain with the help of a long flexible linker. Since mutations in CTD domain impede interference mechanism and CTD undergoes conformational changes (2), we speculate that the linker region allows CTD to move towards and away from the helicase domain (indicated by a double-headed arrow). This is likely to facilitate the bolting of Cas3 onto the non-target strand. Upon recruitment, Cas3 HD domain nicks the displaced non-target strand and subsequently ssDNA strand is fed continuously to the nuclease domain via a directed channel (indicated by a pale brown cylinder) formed at the interface between helicase and CTD domains.

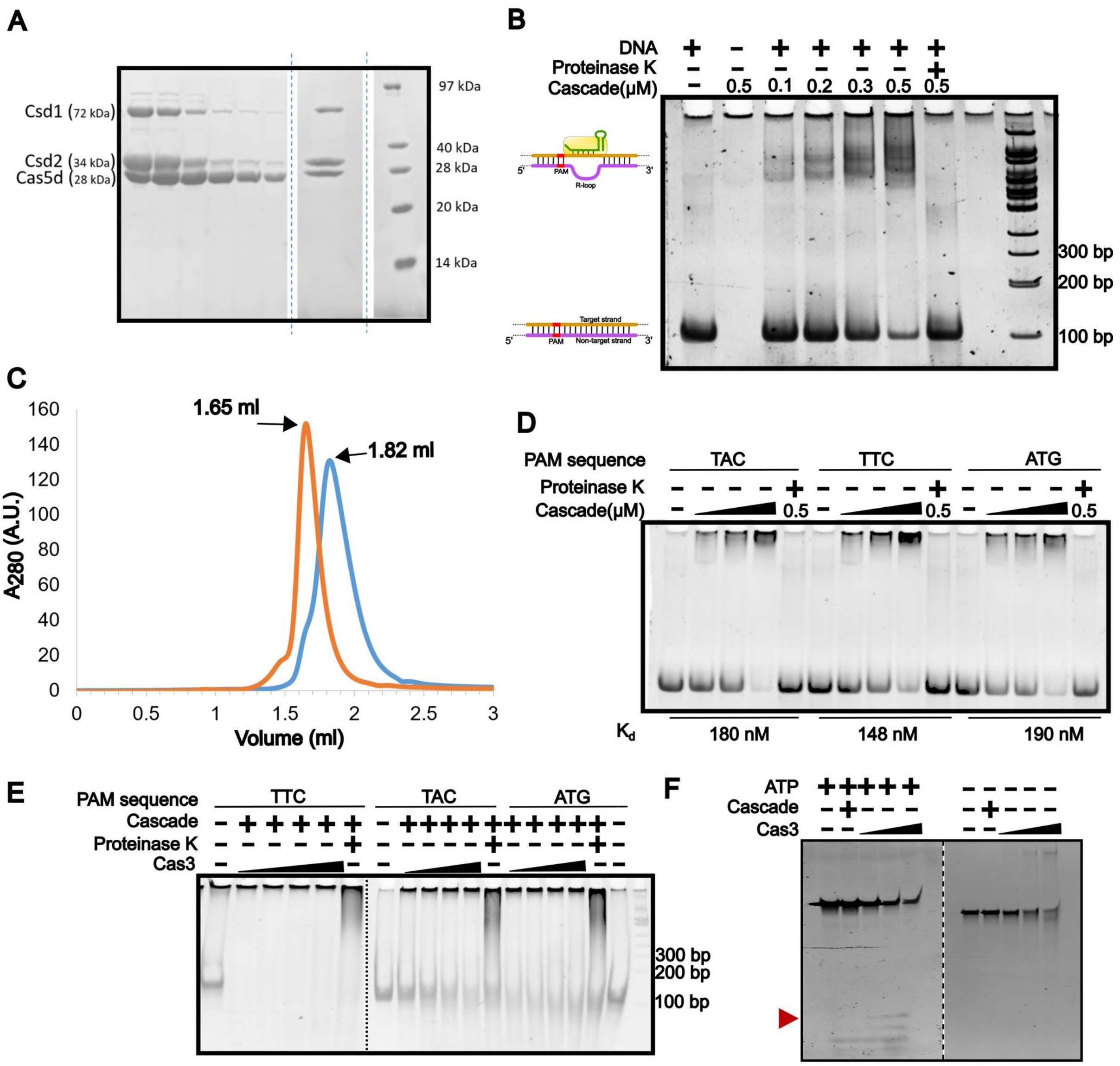


**Figure S5: Characterization of target DNA interaction with Cascade/I-C and Cas3/I-C,** Related to Figure 3, 4 and 5

(A) Purification of the Cascade/I-C complex was achieved by co-expressing Cas proteins and CRISPR array. Csd1 (Cas8c), Cas2 (Cas7) and Cas5d (Cas5) can be observed in SDS-PAGE which shows the affinity-purified (lanes 1-6) and SEC-purified fractions of Cascade/I-C complex (lane 7). The dotted line indicates a discontinuity in the gel for the purpose of clarity.

(B) A 100 bp target DNA that is complementary to crRNA was designed with a 6-FAM label at the 5’end of the non-target strand. In the presence of an increasing concentration of Cascade/I-C, a change in the migration of DNA was observed. Bound DNA fraction was released after the addition of proteinase K, which confirms the formation of the target DNA-Cascade/I-C complex.

(C) DNA bound and free Cascade/I-C showed a significant difference in mobility in analytical size exclusion chromatography. Blue profile corresponds to free Cascade/I-C and orange profile corresponds to DNA bound Cascade/I-C.

(D) Target DNAs with varied PAM sequence (TAC, TTC, and ATG) showed an altered binding affinity towards Cascade/I-C (0-500 nM). However, the canonical PAM sequence (TTC) showed a stronger affinity ($K_d$=148 nM) than other sequences.

(E) Interference was checked on target DNA with varying PAM sequences. Target DNA with TTC containing PAM was cleaved by Cas3/I-C (0-500 nM), whereas TAC and ATG were non-functional. The dotted line indicates two separate gels, which are merged for comparison. Cascade/I-C concentration used was 300 nM.

(F) 100 bp Bubble DNA was incubated with an increasing concentration of Cas3/I-C (0-500 nM) in the presence and absence of 1mM ATP. The cleavage products are indicated with a red triangle. In the absence of ATP, we observed a shift in DNA migration which is absent when ATP is added to the reaction. The dotted line indicates two separate gels, which are merged for comparison.

Note: Samples in supplementary figures S5B, S5D, S5E, and S5F were visualized on 15 % native PAGE.

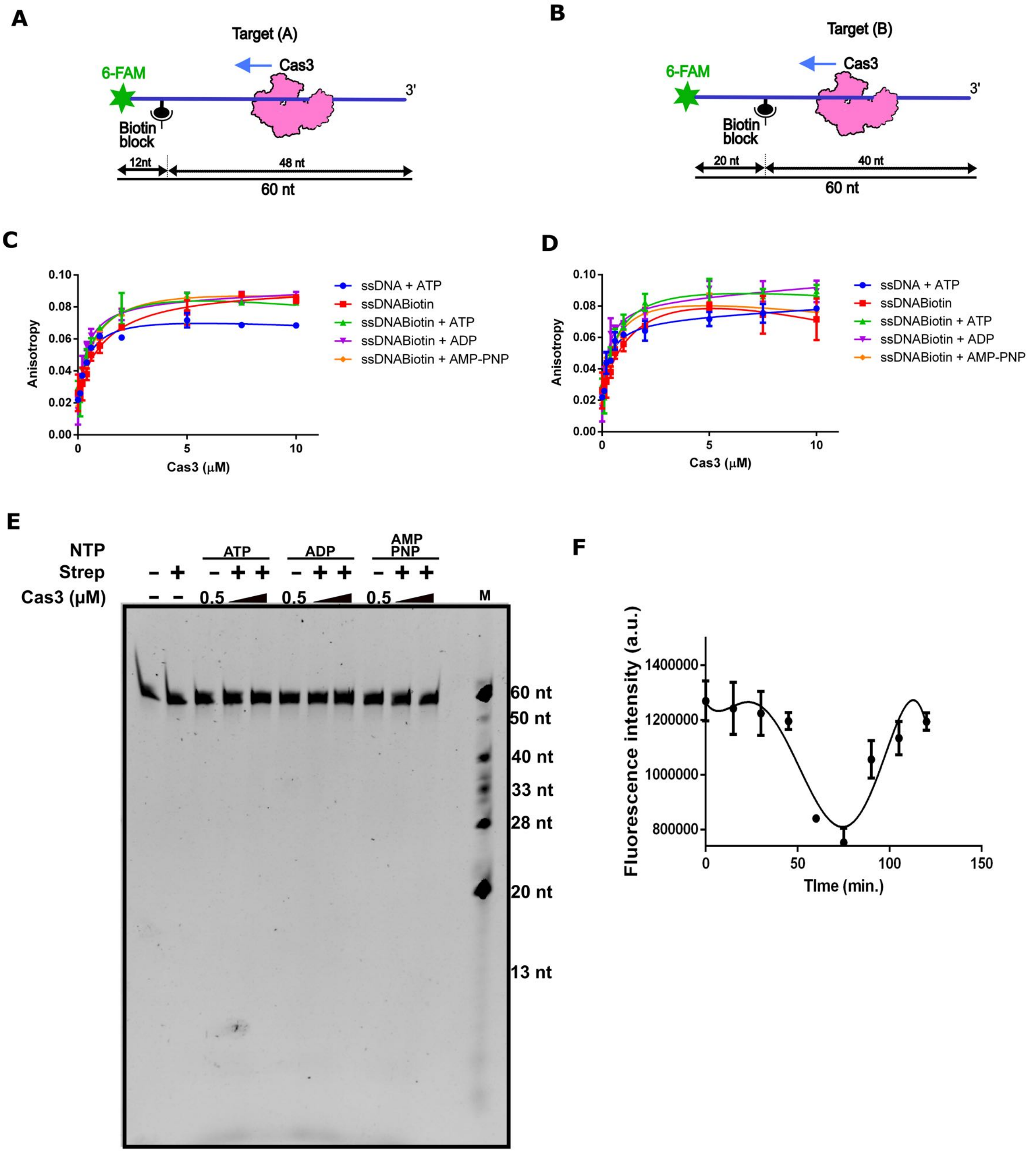


**Figure S6: Roadblock in the translocation of Cas3/I-C stimulates cleavage**, related to figure 4 and 6

(A) & (B) Schematic representation of target DNA (100 bp) as mentioned in Figure 4.

(C) & (D) A roadblock was created using biotin-streptavidin conjugate. An increase in anisotropy was observed when ssDNA mentioned above was incubated with increasing Cas3/I-C concentration, which suggests that Cas3/I-C binds to ssDNA irrespective of the nucleotide.

(E) ssDNA without a biotin block was incubated with Cas3/I-C (250 nM and 500 nM) and 1 mM ATP/ADP/AMPPNP for 30 mins. No cleavage was observed suggesting that in the absence of a blockade Cas3/I-C translocate without exhibiting apparent nuclease activity. Samples were visualized on 20% denaturing PAGE.

(F) DNA construct showed in Figure 4A was incubated with 500 nM of Cas3/I-C and fluorescence intensity was measured over a period of 120 mins. As expected from the reeling mechanism, Cas3/I-

C fluorescence intensity showed a sharp decline for 60 minutes due to quenching of 6-FAM by Iowa Black FQ. Subsequently, there was an increase in fluorescence intensity, suggesting cleavage and release of NTS.

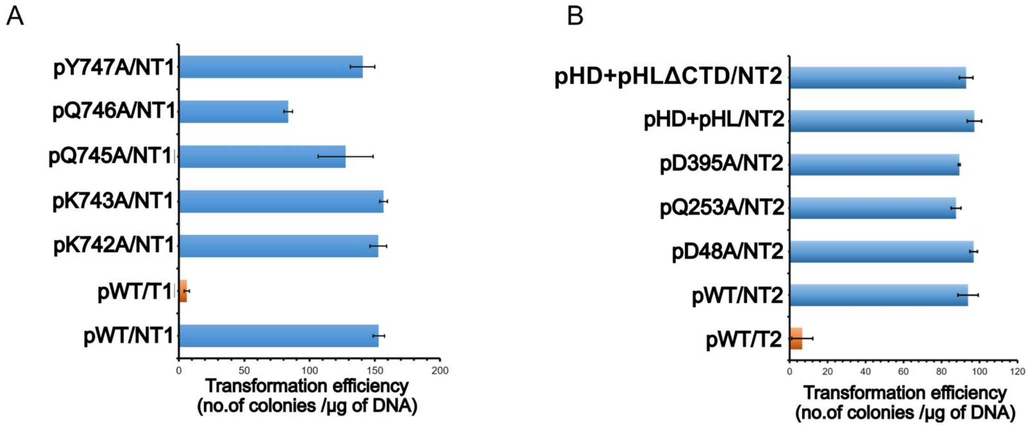


**Figure S7: CRISPR interference against non-targeting plasmid *in vivo*,** Related to Figure 2 and 7

(A) *E. coli* IC-1 was used as a surrogate for Cas3/I-C interference assay. The target sequence was inserted in pUC19 vector (T1), whereas an empty pUC19 vector was used as non-target (NT1). Cas3/I-C and crRNA were expressed through compatible IPTG inducible vectors. High transformation efficiency indicated non-functional CRISPR interference. In the presence of Cas3/I-C (pWT), T1 (shown as an orange bar) showed a reduction in transformation efficiency whereas NT1 showed higher transformation efficiency. Similarly, CTD mutants (pK742A, pK743A, pQ745A, pQ746A, and pY747A) were unable to target NT1.

(B) *E. coli* IC-2 harbouring a spacer that targets pRSF-11 plasmid (T2) was used for *in vivo* studies. Plasmid pST-KT, lacking spacer sequence was used as a non-target plasmid (NT2). Cas3/I-C and its variants were expressed using IPTG inducible pQE2 plasmid and subsequently, transformation efficiency was estimated. Cas3/I-C (pWT), HD nuclease mutant (pD48A), helicase mutant (pQ253A, and pD395A), nuclease-helicase split (pHD + pHL) and nuclease-helicase split without CTD (pHD + pHLΔCTD) were all found to be inactive against NT2. However, as anticipated, Cas3/I-C (pWT) was active against T2 (shown as an orange bar).

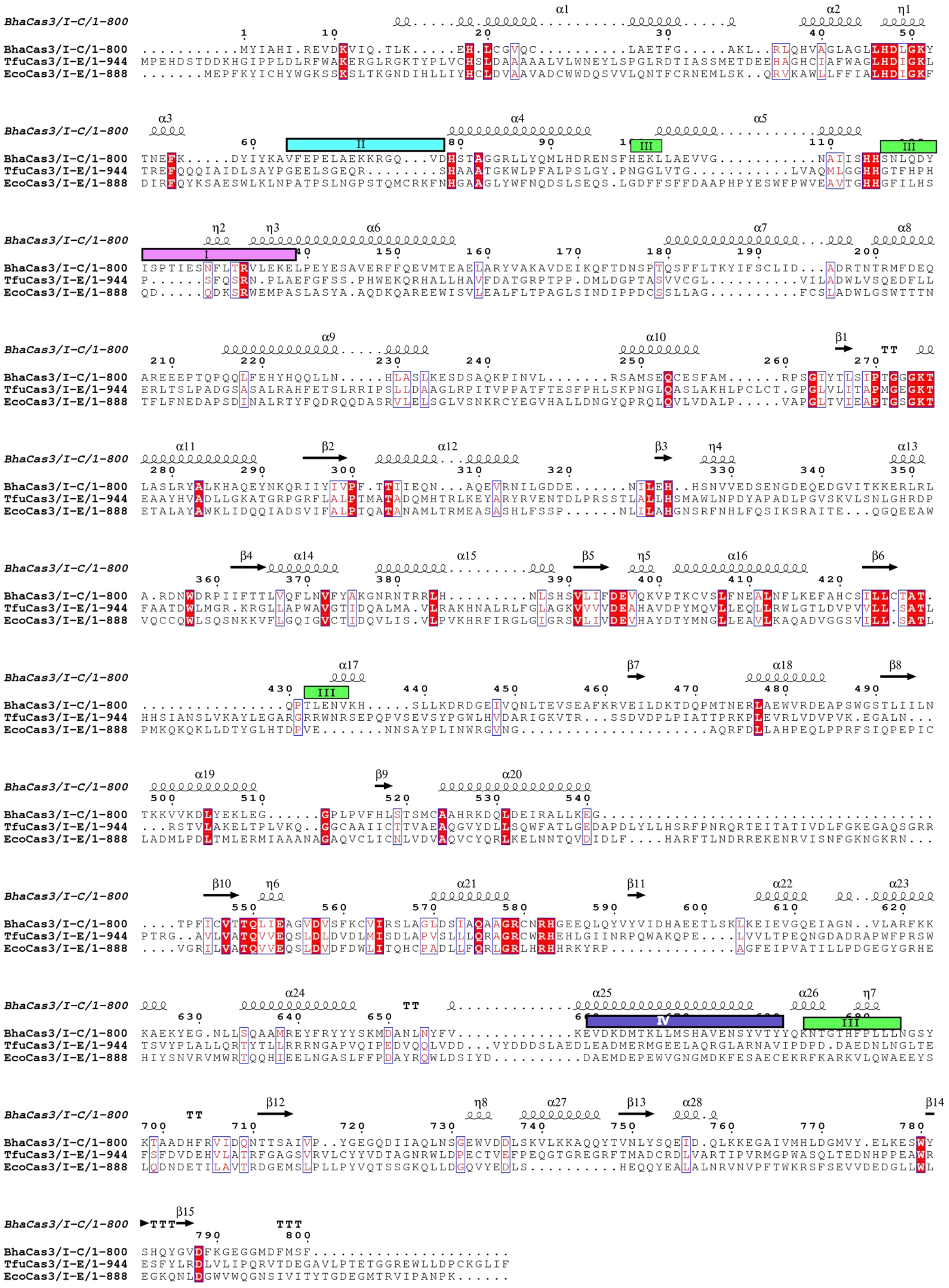

**Figure S8: Sequence conservation among Cas3 from type I-C and type I-E,** related to figure 7
Multiple sequence alignment of BhaCas3/I-C from *B. halodurans* with Cas3 from type I-E shows conservation in the functional motifs. Amino acids showing complete conservation are highlighted in red colour whereas partial conservation is depicted in red letter and boxed. Cas3-Cascade interfaces (I-IV) are shown in coloured bars above the alignment.

### Table S1: Oligonucleotide sequences used in this study

| Oligo name | Sequence (5' to 3') | Description |
|---|---|---|
| Cas3-pQE2-FP | ATGCCATATGATGTACATTGCCCATATTCGA | Amplification of gene encoding Cas3 from *B. halodurans* with restriction sites NdeI and KpnI for pQE2 plasmid. |
| Cas3-pQE2-RP | GGTACCTTAAAACGACATAAAGTCCAT | |
| Cas3-LIC-FP | TACTTCCAATCCAATGCAATGTACATTGCCCATATTCGA | Amplification of gene encoding Cas3 from *B. halodurans* with LIC sites for p1R plasmid. |
| Cas3-LIC-RP | TTATCCACTTCCAATGTTATTATTAAAACGACATAAAGTCCAT | |
| Cas3-D48A | GTGTATTTCCCTAGAGCGTGGAGGAGCCCCG | Along with Cas3-pQE2 and Cas3-LIC primers, these primers were used for PCR based site-directed mutagenesis. |
| Cas3-D395A | GCCATTCCGTCTTAATTTTTGCTGAAGTGCAGAAAGTACCG | |
| Cas3-Q253A | TGGCAAATGACTCGCACGCCTCAGACATAGCAGACC | |
| Cas3-K742A | GACGACTTATCAAAGGTGTTGGCAAAGGCGCAGCAGTATACAG | |
| Cas3-K743A | GTTGACGACTTATCAAAGGTGTTGAAAGCGGCGCAGCAGTATA | |
| Cas3-Q745A | CAAAGGTGTTGAAAAAGGCGGCGCAGTATACAGTCAACCTTT | |
| Cas3-Q746A | AAGGTGTTGAAAAAGGCGCAGGCGTATACAGTCAACCTTTATTC | |
| Cas3-Y747A | GTGTTGAAAAAGGCGCAGCAGGCTACAGTCAACCTTTATTCACA | |
| Cas3'-FP | GGAATTCCATATGGCTATGTCTGAGCAGTGCGAGTCATTT | Amplification of helicase and C-terminal domain of Cas3 from *B. halodurans* (Amino acid 249-800)with restriction sites NdeI and KpnI for insertion in pQE2 plasmid |
| Cas3"-RP | ATACGGGGTACCTTAAGACCGCAGTACATTAATTGGTTTCTGCGC | Amplification of nuclease domain of Cas3 from *B. halodurans* (Amino acid 1-248) with restriction sites NdeI and KpnI for insertion in pQE2 plasmid. |
| Δ*hns*-FP | ATTATTACCTCAACAAACCACCCCAATATAAGTTTGAGATTACTA CAATGTATGAATATCCTCCTTAGTT | For deletion of *hns* gene from Δcas3 strain of *E. coli K-12* |
| Δ*hns*-RP | GATTTTAAGCAAGTGCAATCTACAAAAGATTATTGCTTGATCAGG AAATCTGTAGGCTGGAGCTGCTTCG | |
| Cas5-FP | TACTTCCAATCCAATGCAATGAGAAACGAAGTCCAATTTGAGCTA TTT | Amplification of *cas5*, *cas8c* and *cas7* as a single construct from *B. halodurans* with LIC site for p1R |
| Cas7-RP | TTATCCACTTCCAATGTTATTATTACTGGCCATCAATCACTT | |
| Δ*cas3*-FP | AGCCCGCTGATATCATCGATAATACTAAAAAAACAGGGAGGCTA TTAATGGGCGCGCCTACCTGTGACGG | For deletion of *cas3* (*ygcB*) gene from *E. coli K-12* |
| Δ*cas3*-RP | ATCGTCATTGATAACAATCATTCCCGAAGTTATTTGGGATTTGCA GGGATAACTTCATTTAAATGGCGCG | |
| F1-T7-FP | ATTCGAGCTCGGTACCCGGGGATCCTAATACGACTCACTATAGGG GAATTGTGAGCGGATAACAATTCCCCTCTAGAAATAATTTTGTT | For integration of *cascade/I-C*, using clonetegration into P21 *attB* site of IYB5101. C1-T7-FP was used to insert T7 promoter upstream of *cascade/I-C*. Other oligonucleotides were used to amplify *cascade* from *B. halodurans* |
| F2-*cas5* | AGCTCAAATTGGACTTCGTTTCTCATGGTATATCTCCTTCTTAAAG TTAAACAAAATTATTTCTAGAGGGGAATTGTTATCCGCTCACA | |
| F3-*cas5* | TTTAACTTTAAGAAGGAGATATACCATGAGAAACGAAGTCCAAT TTGAGCT | |
| F4-*cas7*-RP | CATGCATCTCGAGGCATGCCTGCAGTTACTGGCCATCAATCACTT CAACA | |
| Helicase-act-long | AGTGACTTCTGAGGTGTAGGTTACTCACGCTGCTAAGAATGTAGT TTAAGGTGCGTGAGTCCGTGTGTAG | For helicase activity assays. Helicase-act-long was annealed to either of the two 6-FAM labelled |
| Helicase-act-3'short | TTAGCAGCGTGAGTAACCTACACCTCAGAAGTCACT-6-FAM3' | |

| | | |
|---|---|---|
| Helicase-act-5'short | 5'-6-FAM-CTACACACGGACTCACGCACCTTAAACTACATTC | oligonucleotides to form 3' or 5' overhang. |
| F5-Target-TTC-FP | AAACAGCTATGACCATGATTACGCCAAGCTTTTCTAACTCCATCCATAACTAAGCAACCT | These oligonucleotides have 3' complementarity. 100 bp fragment was generated using these oligonucleotides in a PCR reaction and inserted in pUC19 vector to generate Target plasmid (T1) |
| F6-Target-RP | CAGTGAATTCGAGCTCGGTACCCGGGGATCCTACTAGTACAGGTTGCTTAGTTATGGATG | |
| Target-TTC-FP | TACTTCCAATCCAATGCAGGTACCATCGAAAGTAACTCCATCCATAACTAAGCAACCTGT | These oligonucleotides have 3' complementarity. 100 bp DNA construct was generated using PCR |
| Target-TAC-FP | TACTTCCAATCCAATGCAGGTACCATCGAATGTAACTCCATCCATAACTAAGCAACCTGT | |
| Target-ATG-FP | TACTTCCAATCCAATGCAGGTACCATCGATACTAACTCCATCCATAACTAAGCAACCTGT | |
| Targert-RP | TAATAACATTGGAAGTGGATAAAAGCTTTCGATTACTAGTACAGGTTGCTTAGTTATGGA | |
| Dloop-AAG-FP | TACTTCCAATCCAATGCAGGTACCATCGAAAGTAACTCCATCCATAACTAAGCAACCTGTACTAGTAATCGAAAGCTTTTATGGACTTCCAATGTTATTA | These oligonucleotides were annealed to generate 100 bp construct for DNA loop related experiment. (Figure 3c and 3d) |
| Dloop-TTC-RP | TAATAACATTGGAAGTCCATAAAAGCTTTCGAAATGATCATGTCCAACGAATCAATACCTACCTCAATCTTTCGATGGTACCTGCATTGGATTGGAAGTA | |
| ssDNA-biotin_12 | ATGAAGGTT-Biotin-AGGTATACTCCATGCCTAGGTTCATTGAGGTAGGTATTGATTCGTTGGACA-6-FAM-3' | ssDNA target with biotin at 12$^{th}$ position |
| ssDNA | ATGAAGGTTAGGTATACTCCATGCCTAGGTTCATTGAGGTAGGTATTGATTCGTTGGACA-6-FAM-3' | ssDNA target without biotin |
| ssDNA-biotin_20 | ATGAAGGTTAGGTATACTCC-Biotin-ATGCCTAGGTTCATTGAGGTAGGTATTGATTCGTTGGACA-6-FAM-3' | ssDNA target with biotin at 20$^{th}$ position |

### Table S2: List of strains used in this study

| Strain | Genotype | Source |
|---|---|---|
| *E. coli* BW25113 | Δ(*araD-araB*)567, Δ*lacZ4787*(::rrnB-3), λ-, rph-1, Δ(*rhaD-rhaB*)568, *hsdR514* | CGSC#: 7636 |
| *E. coli* IYB5101 (Wt) | F-Δ(*araD-araB*)567 Δ*lacZ4787* (::rrnB-3) λ- rph-1 Δ(*rhaD-rhaB*)568 *hsdR514 araB*::T7-RNAp-*tetA*, TetR | Kind gift from Prof. Udi Qimron |
| *E. coli* JW1225-2 | F-, Δ(*araD-araB*)567, Δl*acZ4787*(::rrnB-3), λ-, *Δhns-746::kan*, rph-1, Δ(*rhaD-rhaB*)568*, hsdR514,* Kan$^R$ | CGSC#: 9111 |
| *E. coli* BL21-AI | F- *ompT hsdsB*(rB-, mB-) *gal dcm araB*:: T7-RNAp-*tetA*, Tet$^R$ | Invitrogen |
| *E. coli* BL21(DE3) | F- *ompT hsdsB*(rB-, mB-) *gal dcm* λ(DE3) | |
| *E. coli* TOP10 | F- *mcrA* Δ(*mrr-hsdRMS-mcrBC*) ϕ80*lac*ZΔM15 Δ *lac*X74 *recA1 araD139* Δ(*araleu*)7697 *galU galK rpsL endA1 nupG*, Str$^R$ | Invitrogen |
| *E. coli* IC-1 | IYB5101 *Δcse1-cas6* P21::*cas5-cas7*, Cam$^R$ | This study |
| E. coli IC-2 | JW1225-2 *Δhns Δcas3*, Cam$^R$ | This study |

### Table S3: List of plasmids used in this study

| Plasmid name | Description | Source |
|---|---|---|
| pKD46 | *ori* R101, *repA101ts*, $Amp^R$, *araC*, expresses λ Red genes (*gam-bet-exo*) under the control of arabinose inducible promoter ($P_{araBAD}$). | CGSC #7739 |
| pOSIP-CT | *ori* Rγ, *ori* pUC, $Cam^R$, *attP* P21, *ccdB*, λ (*cI857*) encodes P21 integrase under the control of λ promoter (λ pR). | Addgene #45981 |
| pUC19 / pNT1 | *ori* PBR322, $Amp^R$, for gene insertion under the control of lac promoter. | New England Biolabs |
| pQE2 | *ori* ColE1, $Amp^R$, expresses gene of interest to synthesize N-terminal 6xHis tagged protein under the IPTG inducible T5 promoter. | |
| pST-KT / pNT2 | *ori* PBR322, $Kan^R$ | Kind gift from Dr. VK Nandicoori |
| pRSF-11 / pT2 | *ori* RSF, $Kan^R$, cloning and expression vector with T7 inducible promoter | |
| pET StrepII TEV LIC cloning vector (p1R) | *ori* pMB1, $Kan^R$*, lacI*, expresses the gene of interest to synthesize N-terminal StrepII tagged protein under the control IPTG inducible promoter (PT7lac). | Addgene #29664 (Scott Gradia) |
| pET StrepII TEV co-transformation cloning vector (p13SR) | *ori* CloDF13, $Spc^R$, *lacI*, expresses the gene of interest to synthesize N-terminal StrepII tagged protein under the control IPTG inducible promoter (PT7lac). | Addgene #48328 (Scott Gradia) |
| pWT-1 | p1R inserted with *cas*3 gene encoding N-terminal Strep-II tagged protein. | This study |
| pWT-2 | pQE2 inserted with *cas3* gene encoding N-terminal 6xHis tagged protein. | This study |
| pHD | p1R inserted with HD domain (nuclease) of Cas3 encoding N-terminal Strep-II tagged protein. | This study |
| pHL | p1R inserted with DExD/H domain (helicase) of Cas3 encoding N-terminal Strep-II tagged protein. | This study |
| pCas3ΔCTD | p1R inserted with Nuclease and Helicase domain of Cas3 without CTD, encoding N-terminal Strep-II tagged protein. | This study |
| pCascade/I-C | p1R inserted with Cascade proteins (Cas5, Cas8 and Cas7) encoding N-terminal Strep-II tagged protein. | This study |
| pK742A | p1R inserted Cas3 with alanine mutation at K742 amino acid, encoding N-terminal Strep-II tagged protein. | This study |
| pK743A | p1R inserted Cas3 with alanine mutation at K743 amino acid, encoding N-terminal Strep-II tagged protein. | This study |
| pQ745A | p1R inserted Cas3 with alanine mutation at Q745 amino acid, encoding N-terminal Strep-II tagged protein. | This study |
| pQ746A | p1R inserted Cas3 with alanine mutation at Q746 amino acid, encoding N-terminal Strep-II tagged protein. | This study |
| pY747A | p1R inserted Cas3 with alanine mutation at Y747 amino acid, encoding N-terminal Strep-II tagged protein. | This study |
| pT1 | Target DNA sequence with TTC PAM inserted in pUC19 vector | This study |
| pCRISPR/I-C | p13SR plasmid inserted with DNA sequence encoding CRISPR array having 7 copies of same repeat-spacer units. | This study |
| pD48A | pQE2 inserted Cas3 with alanine mutation at D48 amino acid, encoding N-terminal 6x His tagged protein. | This study |
| pD253A | pQE2 inserted Cas3 with alanine mutation at D253 amino acid, encoding N-terminal 6x His tagged protein. | This study |
| pD395A | pQE2 inserted Cas3 with alanine mutation at D395 amino acid, encoding N-terminal 6x His tagged protein. | This study |
| pHD + pHL | pQE2 inserted Nuclease and Helicase domain encoding bicistronic cassette. | This study |
| pHD + pHLΔCTD | pQE2 inserted Nuclease and Helicase domain without CTD encoding bicistronic cassette. | This study |
